# Structural basis of HSP90C, a highly active chloroplastic HSP90 chaperone from *A. thaliana*

**DOI:** 10.64898/2026.01.05.697646

**Authors:** Romain La Rocca, Thomas Chenuel, Céline Bergonzi, Alexandre Maes, Alexandre Pozza, Philippe Meyer

## Abstract

Chloroplasts are the main energy organelles in plants, primary through photosynthesis. Thereby, they are responsible for CO_2_ fixation and dioxygen production, which are essential for living species on Earth. To ensure these processes, numerous proteins encoded from the nuclear DNA need to be imported inside the chloroplast, and eventually to the thylakoids. Whereas the translocation systems from both chloroplastic and thylakoids membranes have been studied in recent years, the stromal route between these two membranes is largely unknown. Notably, the chloroplastic HSP90 (HSP90C) is likely to play an important role in this process, but its structure and molecular mechanisms remain to be unveiled. In this study, we used a combination of structural and biophysical approaches to elucidate the features of *Arabidopsis thaliana*’s HSP90C. Principally, we found that HSP90C has a remarkably high ATPase activity among the HSP90 family proteins. Further investigation allowed us to pinpoint atypical mechanisms responsible for this high activity. First, the N-terminal cap is involved in a disulfide bond that accelerates the ATPase activity of HSP90C. Second, its C-terminal domain features an extension that is mandatory for its dimerization. Third, our crystal structures reveal a wide opening of the HSP90C’s dimer with reduced intermonomeric interfaces. Lastly, we identified a helical switch which is required for HSP90C’s high activity. Three of these four features are due to sequence signatures of HSP90C, which we found to be shared by most of green plants representatives. Our study provides first insights of HSP90C’s non-canonical mechanisms, which will help in the understanding of processes related to protein import in the chloroplast.

## Introduction

Chloroplasts are the energetic cores of plants, being essential for their living but also for humans. Indeed, chloroplasts allow plants to fix CO_2_ in the atmosphere and generate dioxygen through photosynthesis. This unique process occurs in thylakoids, where many proteins work in coordination to ensure energy production. However, most of the genes encoding these proteins are located in the nucleus ^1^. It implies that they have to transit through the chloroplastic and thylakoid membranes, after which they can be imported into the thylakoids. These two crucial steps are performed by specific translocon machineries, namely the TIC/TOC complex for the chloroplastic membrane ^2^, and three different systems for the thylakoid membrane: SEC, Tat and SRP pathways ^3^. One less known step is the route between these two membranes inside the stroma, where some proteins stay in or continue their path to one of the translocon from the thylakoids. Among the proteins that are likely to be involved in this route, HSP90C has been described as an important actor.

HSP90C is a chaperone protein belonging to the HSP90 family. Its gene has been identified initially in 1996 ^4^, and investigated in different studies the years after. Indeed, HSP90C is localized in the stroma, and is essential for the photomorphogenesis of *A. Thaliana* ^5^. Later, HSP90C was identified as an essential factor for thylakoid formation and plant development ^6^. Besides, several studies raised evidence about its role in the stroma. Inoue and co-workers demonstrated that HSP90C is associated with components from the TIC/TOC translocation system, namely Tic40 and Tic110 ^7^. The authors also showed that by using radicicol, which is a specific inhibitor of HSP90, the protein import was greatly decreased. In parallel, other studies identified HSP90C as a partner of thylakoid membrane translocon ^8–10^. Indeed, HSP90C can associate with components of the SEC translocon ^10^, as well with PsbO1, which is a subunit of the PSII ^9^. Most importantly, HSP90C prevents the accumulation of PsbO1 in the stroma, and facilitates its delivery to the thylakoids. Further evidence of a direct link between proteins from the thylakoids and HSP90C has been provided in two studies ^10,11^ in which the thylakoids membrane association factor VIPP1 was found to interact with HSP90C. Thus, HSP90C is believed to guide client proteins from the chloroplast translocon to the thylakoids, as it is located near both these membranes. Recently, the C-terminal tail of HSP90C has been identified as an important region for both HSP90C’s ATPase and client binding ^12^. Besides this study, the mechanism by which HSP90C acts is still poorly known.

To provide insights of its molecular behavior, we investigated HSP90C from *A. Thaliana* at the molecular-scale by means of biochemical and biophysical tools. Using molecular biology constructs, we studied its ATPase activity, refolding activity and dimerization, while investigating its structure with crystallography experiments.

## Results

### HSP90C differs by its sequence and ATPase activity

The HSP90C sequence is mainly characterized by its three domains that are commonly shared among the HSP90 family (Fig. 1A) ^13^. The N-terminal domain (NTD) is responsible for the ATPase binding, which is mandatory for the function of HSP90 proteins ^14^. Following it, the middle domain (MD) triggers the ATP hydrolysis by binding the γ-phosphate of the ATP located in the NTD. The MD is also involved in interactions with client proteins and co-chaperons, as previously demonstrated ^15,16^. Finally, the C-terminal domain (CTD) is responsible for the dimerization of HSP90, thereby being a key actor of the HSP90 cycle ^17^.

**Figure 1:**
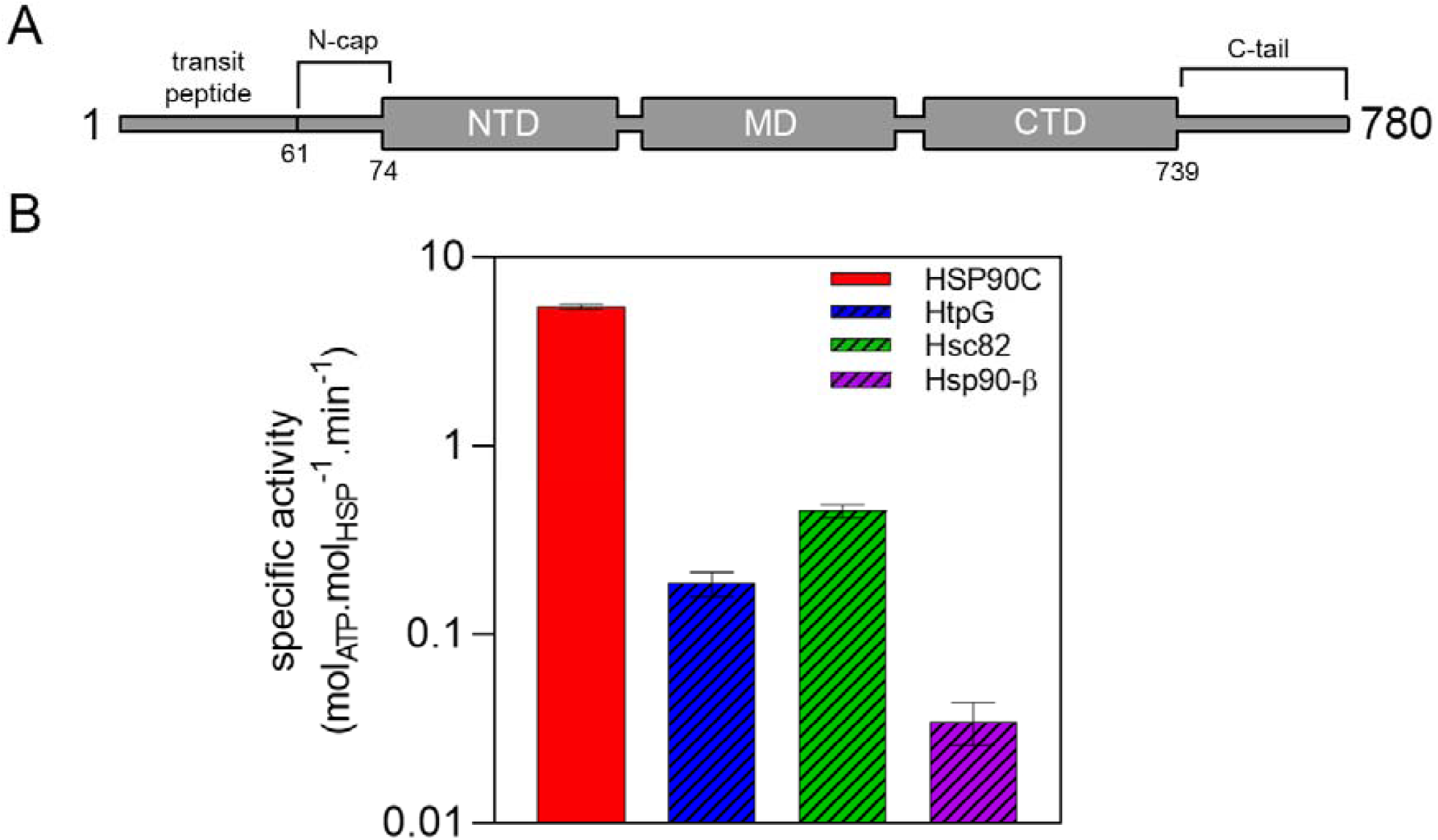
Sequence (A) and ATPase activity test (B) of HSP90C. A: HSP90C includes the three main canonical domains of HSP90 proteins: the N-terminal domain (NTD), the middle domain (MD) and the C-terminal domain (CTD). In addition, it featuresa transit peptide (1-60), a 13 residues-long N-terminal cap (N-cap) and a 41 residues-long C-terminal tail (C-tail). B: HSP90C’s ATPase activity among HSP90 proteins from other species. HtpG is from *E. coli* (bacteria), Hsc82 is from *S. cerevisae* (yeast), and Hsp90-β is from *H. sapiens* (human). Activity is displayed as a log scale.

In addition to the 3 canonical domains of HSP90 proteins, HSP90C contains 2 specific parts that are less conserved among the HSP90 family. The N-terminal cap is the part of the sequence following the transit peptide and preceding the NTD (61-74) (Fig. 1A). In the HSP90 family, this motif is mostly distinguishable by its position rather than its amino-acid composition, which is very different from one HSP90 to one other. For instance, the N-terminal cap of Grp94 (the human endoplasmic reticulum HSP90) is 51 residues long ^18^, whereas the one from TRAP-1 (the human mitochondrial HSP90) contains 26 amino-acids ^19^ However, despite their different nature, the N-terminal caps from both Grp94 and TRAP-1 have been shown to decrease the ATPase activity of the protein, thereby suggesting a conserved role of the N-cap in the HSP90 family ^18,19^. At the very end of the sequence, HSP90C exhibits a C-terminal tail, which is 41 residues long (739-780) (Fig. 1A). Similarly to the N-terminal cap, the C-terminal tail is very variable among all of the HSP90 homologs. In the case of eukaryotic cytosolic HSP90 proteins, this part often shows a MEEVD motif at the very end of the tail, which is responsible for interaction with co-chaperons ^13^. For Grp94, this motif is replaced by a KDEL sequence that allows its retention onto the ER ^20^. In HSP90C, a part of this tail (750-780) is involved in both ATPase activity and client binding ^12^. The role of both N-terminal cap and C-terminal tail of HSP90C has not been fully assigned yet and remains hard to predict, given the high variability of these two sequences in all HSP90 proteins.

Preliminary to our study, we initially conducted an ATPase activity test of HSP90C from *A. thaliana* to evaluate its activity among the HSP90 family (Fig. 1B). We performed the same experiment on 3 homologs from different organisms: HtpG (Bacteria, *E. coli*), Hsc82 (Yeast, *S. cerevisiae*), and Hsp90-β (Human, *H. sapiens*). We observed that HSP90C has a remarkable ATPase activity among the HSP90 family, which is higher from 10-fold (Hsc82) to 100-fold (Hsp90-β) in comparison to the 3 HSP90 homologs that we tested. This result highlights the singularity of HSP90C from *A. thaliana*, which may result from specific molecular mechanisms that remain to be unveiled.

To clarify these mechanisms, we first focused on experiments of deleted forms of the N-terminal cap, in order to assess its impact on HSP90C activity. Second, we used the same approach on the C-terminal tail. Lastly, we studied HSP90C using X-ray crystallography to obtain its structure, and further investigated a sequence signature that we identified thereby.

### The N-terminal cap regulates HSP90C activity through a disulfide bond formation

To assess the role of the N-terminal cap, we cloned and produced 3 constructions of HSP90C (Fig. 2A): the mature WT form of the protein (61-780), one form additionally lacking the 4 first residues of the N-cap (65-780), and a construction lacking the entire cap (74-780). We then measured both the ATPase and refolding activities (Fig. 2B). Given that the small motif at the start of the N-terminal cap contains one cysteine (C61), the presence or the absence of a reducing agent in the reacting medium may be important. Therefore, we conducted both ATPase and refolding activity assays in reducing and non-reducing conditions. We noticed a decreased ATPase activity and an increased folding activity between 61-780 and the two other constructions lacking the CDAA sequence (65-780 and 74-780) in non-reducing conditions. In contrary, we observed that ATPase and refolding activities were very comparable between the 3 constructions in reducing conditions. Then we measured the ATPase activity of the same 3 constructions after the addition of H_2_O_2_ following the reduction of HSP90C (Fig. 2C). The results showed that H_2_O_2_ tends to restore the ATPase activity of 61-780 back to its level before the reduction, but not for the 65-780 and 74-780 constructs. These experiments show that C61 is involved in the regulation of HSP90C ATPase and foldase activities that is most likely to occur via a disulfide bond formation.

**Figure 2:**
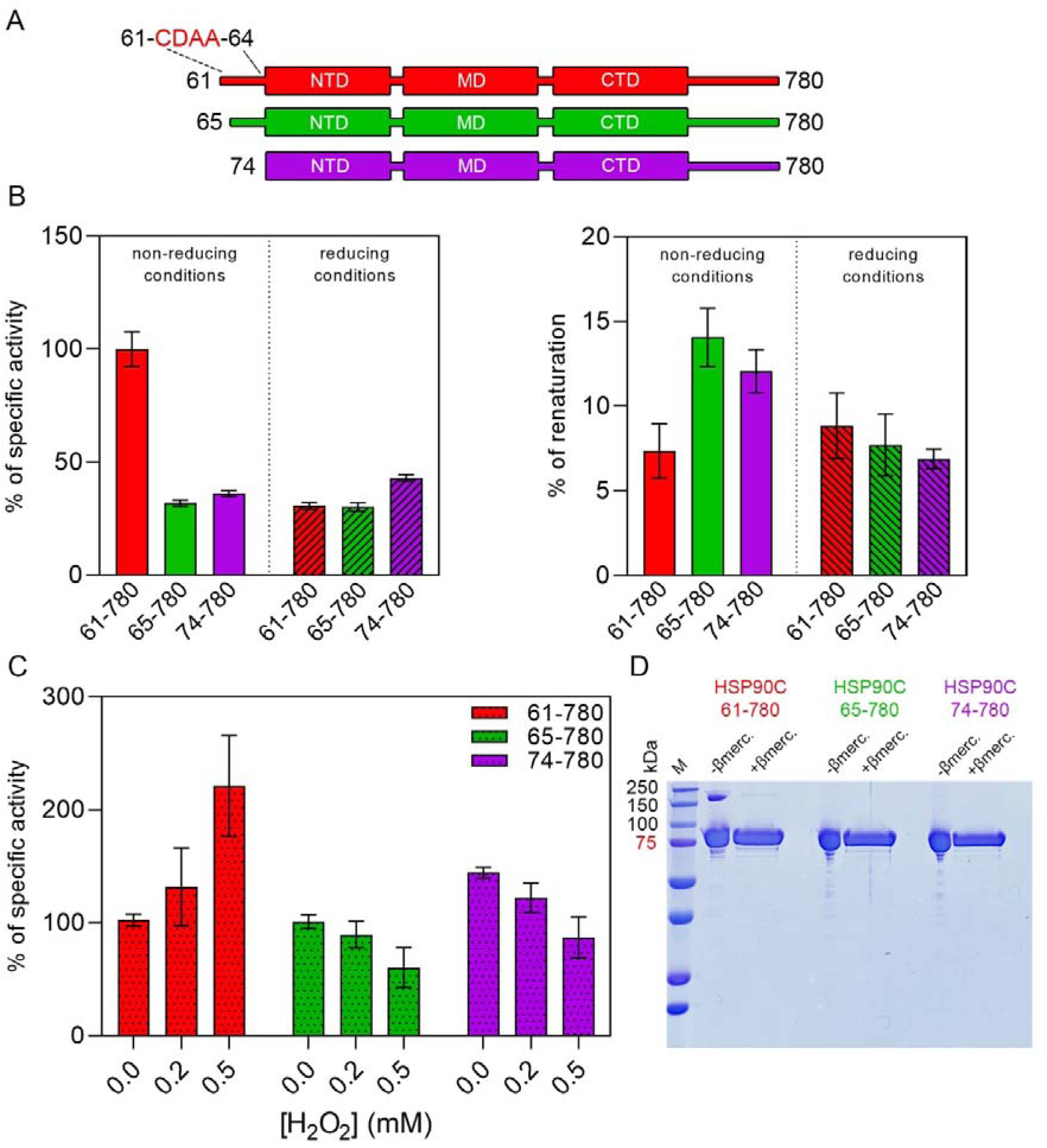
Redox regulation by the N-cap of HSP90C. A: Mature HSP90C construction (red) and N-cap truncated constructions (green and purple) of HSP90C used in this study. B: ATPase activity (left panel) and refolding activity assays (right panel) of HSP90C constructions, in non-reducing and reducing conditions. Both scales are shown as percentages, relative to the mature HSP90C activity (ATPase assays) and to the condition without HSP90C (refolding activity assays). C: ATPase activity tests in the presence of H_2_O_2_. D: SDS-PAGE gel of HSP90C constructions, with or without a reducing agent.

To further investigate this regulation, we conducted an SDS-PAGE gel with or without a reducing agent (Fig. 2D). This experiment revealed a band corresponding to the dimeric form of HSP90C (170 kDa), which is only noticeable for the 61-780 construction without any reducing agent. By adding 2-mercaptoethanol, the dimeric band disappeared. Besides, we saw for all the constructions a band corresponding to the monomeric form of HSP90C (85 kDa), which is not affected by the reducing agent. This suggests that C61 is likely to be involved in an intermonomeric disulfide bond formation, thereby stabilizing the dimeric form of HSP90C. We then tried to identify a putative partner of C61 involved in the disulfide bond formation (Fig. S1). According to the AlphaFold3 structure prediction of HSP90C, most of its cysteines are likely to be located too far from the N-terminal cap (Fig. S1A). Although both C61 appears to be distant in this prediction, they are located in a highly flexible region difficult to predict, which is capping the other HSP90 dimer in both Grp94 and TRAP-1 structures ^18,19^. Thus, C61 is most likely to connect to C61 from the other monomer, or to C165. To confirm or infirm these hypotheses, we cloned and produced a punctual mutant of C165 of HSP90C (C165S, Fig. S1B), and conducted ATPase activity tests on this mutant. The C165S mutant showed a similar sensitivity to H_2_O_2_ in comparison to the WT (61-780) (Fig. S1C). Moreover, SDS-PAGE gel analysis revealed that the C165S mutant is still able to form the dimeric bond without 2-mercaptoethanol (Fig. S1D). These results suggest that C165 is not involved in the disulfide bond formation with C61.

Taken together, these experiments indicate that C61 is involved in an intermonomeric disulfide bond formation, which regulates both its ATPase and refolding activities.

### A C-terminal extension of the HSP90C CTD is mandatory for its dimerization

After investigating the role of the N-terminal cap of HSP90C in its function, we conducted the same experiments on C-terminal tail-truncated constructions (Fig. 3A). We cloned and produced a construction containing the entire tail (74-780), one other lacking most of the tail (74-745), and a last one lacking the entire tail (74-739). The ATPase and refolding activity assays on these constructions revealed that most of the activity of HSP90C was conserved after the truncation of the major part of the C-tail (745-780) (Fig. 3B). However, when the 739-745 region is lacking (RWGRVE), both ATPase and refolding activities were dramatically lower. As this hexapeptide is very close to the CTD which is responsible for the dimerization of HSP90C, we performed SEC-MALS analysis to evaluate the ability to dimerize of these 3 constructions (Fig. 3C). The SEC-MALS data revealed that the 74-745 and 74-780 constructions have a molecular weight corresponding to the dimeric form of HSP90C (150 kDa). Moreover, a shift of the elution volume between these 2 constructions suggests that the shape of the protein may be different. For the 74-739 construct, we observed a major peak corresponding to the monomeric form of HSP90C (80 kDa), while the dimeric peak was almost absent. Consequently, the elution volume was greatly different in comparison to 74-745 and 74-780. These experiments show that a small peptide of the C-terminal tail of HSP90C (RWGRVE) is mandatory for its dimerization.

**Figure 3:**
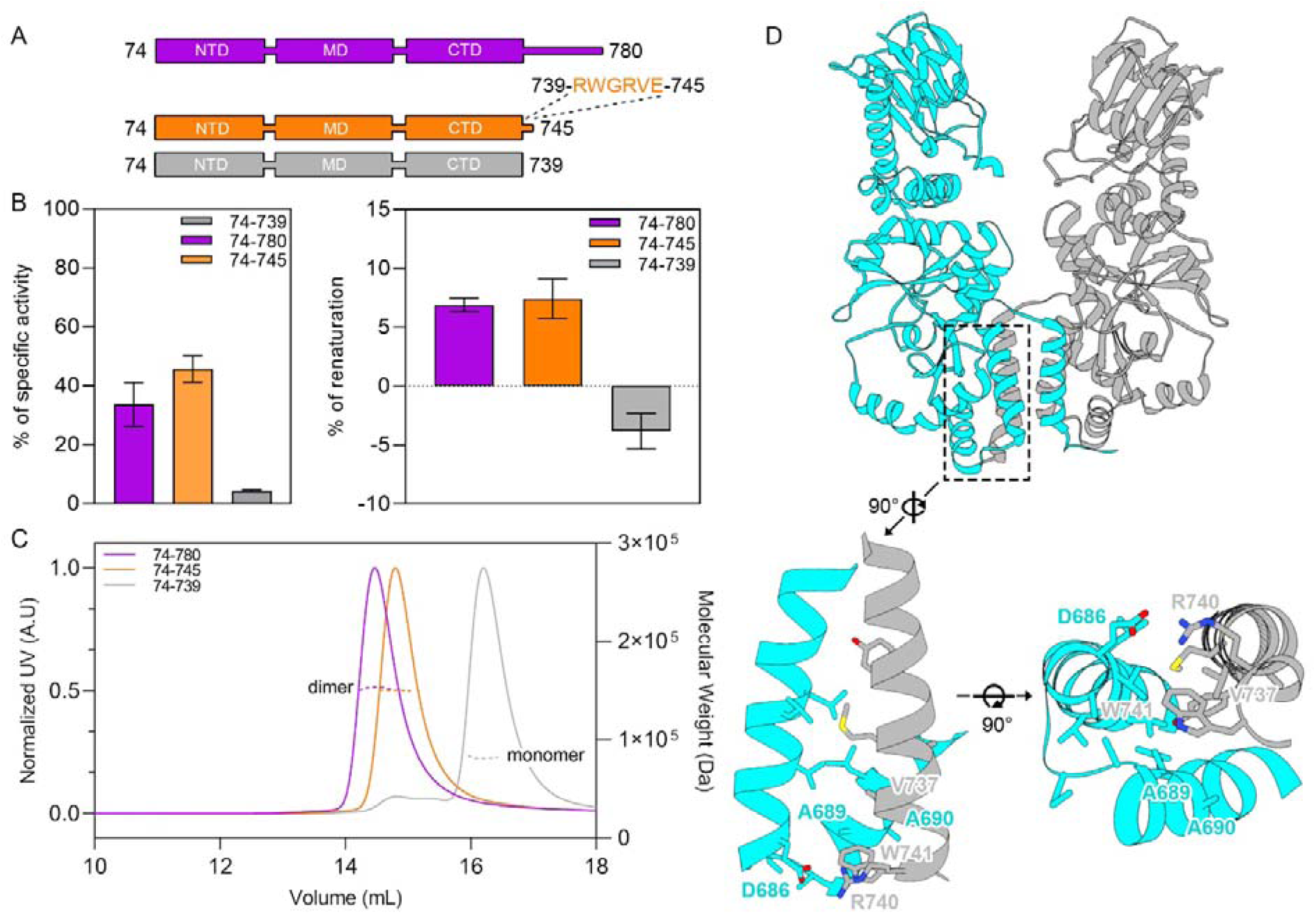
Role of the CTD extension of HSP90C. A: HSP90C construction with full C-tail (purple) and C-tail truncated constructions (orange and grey) of HSP90C used in this study. B: ATPase activity (left panel) and refolding activity assays (right panel) of HSP90C C-tail truncated constructions. C: SEC-MALS experiment with C-tail truncated HSP90C constructions, showing the incapacity of 74-739 to dimerize. D: Crystal structure of the 333-745 HSP90C dimer (PDB: 9SWT). Both monomers are represented with different colors (cyan and grey). The lower panel shows a new dimeric interaction occurring in HSP90C, involving R740 and W741 from the CTD extension.

To further investigate, we crystallized a construction of HSP90C that includes the RWGRVE peptide. This construction contains the MD, the CTD and the RWGRVE peptide of HSP90C (333-745). We obtained the structure of the 333-745 dimer at a resolution of 3.0Å (Fig. 3D). It revealed that the RWGRVE peptide is interacting with the CTD of the other monomer of HSP90C. This interaction consists of a first polar interface between R740 and D686 residues. Following this, the next tryptophan (W741) of the RWGRVE peptide stands at the bottom of a hydrophobic pocket within the helix bundle. Indeed, W741 interacts with V737 from the same monomer, while its faces A689 and A690 from the other monomer. Thereby, this RW motif creates a small interface that stabilizes the dimer.

Further analysis of this structure also allowed us to notice a wide dimer opening among the HSP90 proteins.

### HSP90C dimer has a wide opening among HSP90 family

The crystallographic structure of the 333-745 dimer arbors a wide-opened shape, which is unique among the HSP90 family members. To evaluate this feature, we compared the 333-745 dimeric structure of HSP90C with all previously published structures of MD and CTD dimers from other homologs. These structures are the MD and CTD dimers of Grp94 from canine (PDB: 2O1T), Hsc82 from yeast (PDB: 2CGE), and Hsp90α from human (PDB: 7RY1). Then we performed an analysis of the surfaces using the InterProSurf server (Fig. 4). For each of these 4 structures, the buried surface area per residue (BSAPR) was calculated, which relates the proximity of HSP90 amino-acids to the ones from the other monomer (Fig. 4A). Thereby, four regions were identified as close regions between monomers (numbered from 1 to 4). The results show comparable interfaces in the regions 1, 2 and 4. However, HSP90C has no dimeric interaction in the region 3 (green), unlike all other HSP90 analyzed here (Fig. 4A, 4B). This region corresponds to the middle helices of the HSP90C dimer (650-670), which stands above the CTD dimerization helix bundle (Fig. 4C). Moreover, we superimposed these four HSP90 MD-CTD structures on their CTD on one monomer (Fig. S2). This superimposition allows us to evaluate the relative position of the other HSP90 monomer. In addition to the center helices previously identified (650-670), we observed that the CTD of HSP90C is largely shifted to the outside of the lumen. Particularly, the secondary structures of the C-terminal domain from the other monomer are shifted relative to the other HSP90. Thus, HSP90C’s dimeric conformation is markedly different among the HSP90 family.

**Figure 4:**
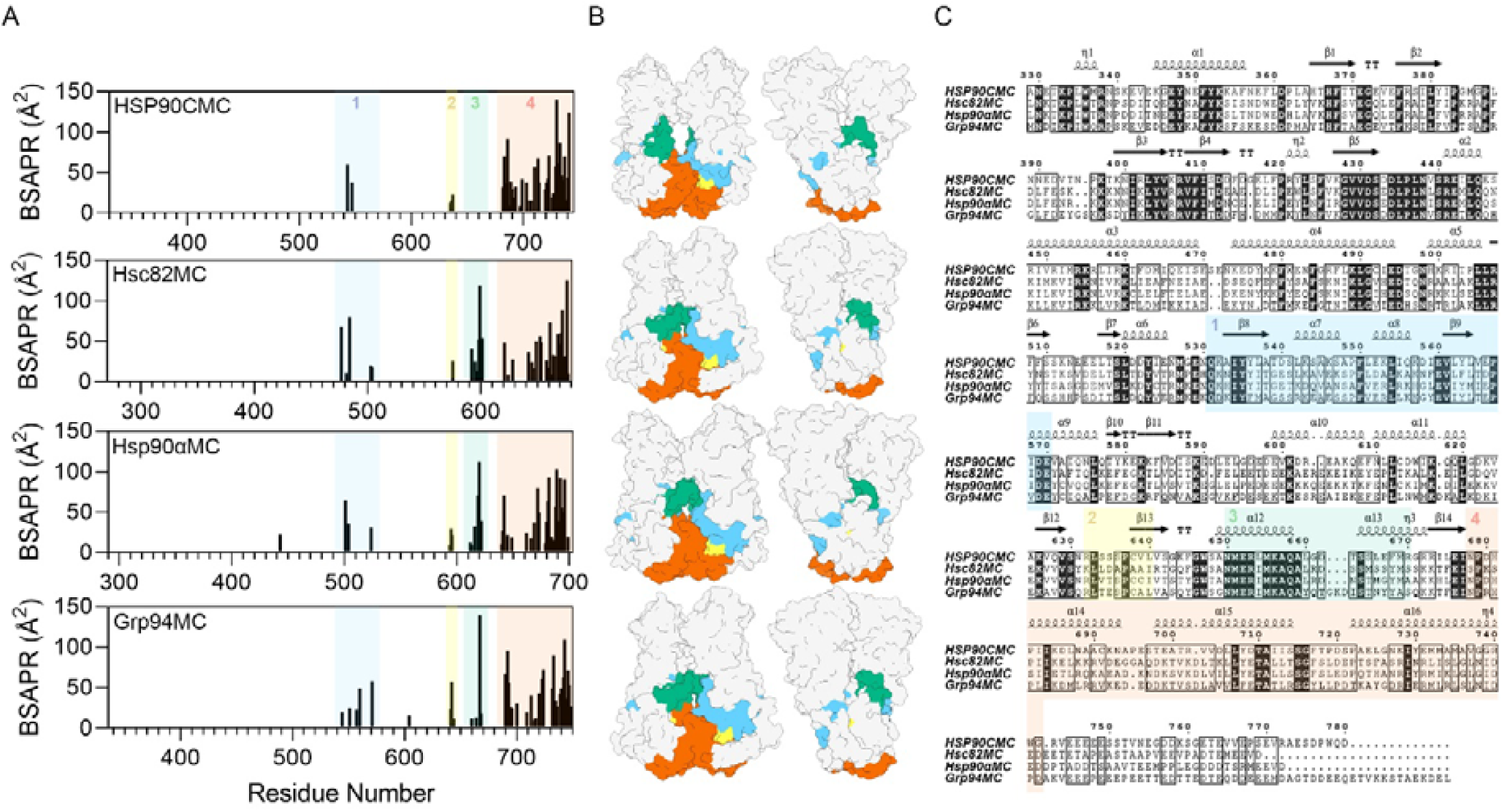
Intermonomeric interfaces between HSP90 monomers of HSP90 MD-CTD dimers from different species. Dimers includes only the middle domains and C-terminal domains of HSP90C from *A. Thaliana* (PDB: 9SWT), Grp94 from *C. lupus familiaris* (PDB: 2O1T), Hsc82 from *S. cerevisiae* (PDB: 2CGE), and Hsp90α from *H. sapiens* (PDB: 7RY1). A: Buried Surface Area Per Residue (BSAPR) for the four HSP90 MD-CTD dimers represented here, calculated with InterProSurf. The X scale (Residue Number) is based on the multiple sequence alignment performed with Clustal Omega. The four main intermonomeric interfaces are represented as blue (1), yellow (2), green (3) and orange (4). B: Main interfaces represented on HSP90C, Hsc82, Hsp90α and Grp94 dimers (from top to bottom). C: ESPript ^42^ representation of the multiple sequence alignment between the four HSP90 dimers represented here, with corresponding secondary structures from the HSP90C dimer. Intermonomeric interfaces are shown as blue (1), yellow (2), green (3) and orange (4).

In a second time, this structure allowed us to identify a conformational switch that appears to be crucial for the activity of HSP90C.

### The crystal structure of HSP90C reveals a unique conformational switch

Subsequently, we obtained the crystal structure of the isolated MD of HSP90C (333-598) at a resolution of 3.0Å, and superimposed both the middle domains from the 333-745 structure and the 333-598 structure (Fig. 5A). The superimposition revealed that most of the domain is very similar between these 2 constructions, except for the α11 helix. In the case of the 333-745 structure, the α11 helix adopts a canonical “curved” conformation, which was previously observed in the MD+CTD structures of Grp94, Hsc82 and Hsp90α homologs. In this case, the α11 helix is separated into 2 parts because of a proline residue that prevents the helix from continuing. Eventually, the proline is preceded by a serine residue which is capping the next part of the helix with a hydrogen bond. For HSP90C, this hydrogen bond does not occur because of the replacement of this conserved serine into an alanine (A548). Nevertheless, the 333-745 construction still adopts this curved conformation. In the case of the 333-598 structure, we observed a unique straight conformation for the α11 helix. This straight conformation is most likely occurring because of the absence of any hydrogen bond stabilizing the second part of the helix. These observations highlight a non-canonical conformation of the α11 helix of HSP90C.

**Figure 5:**
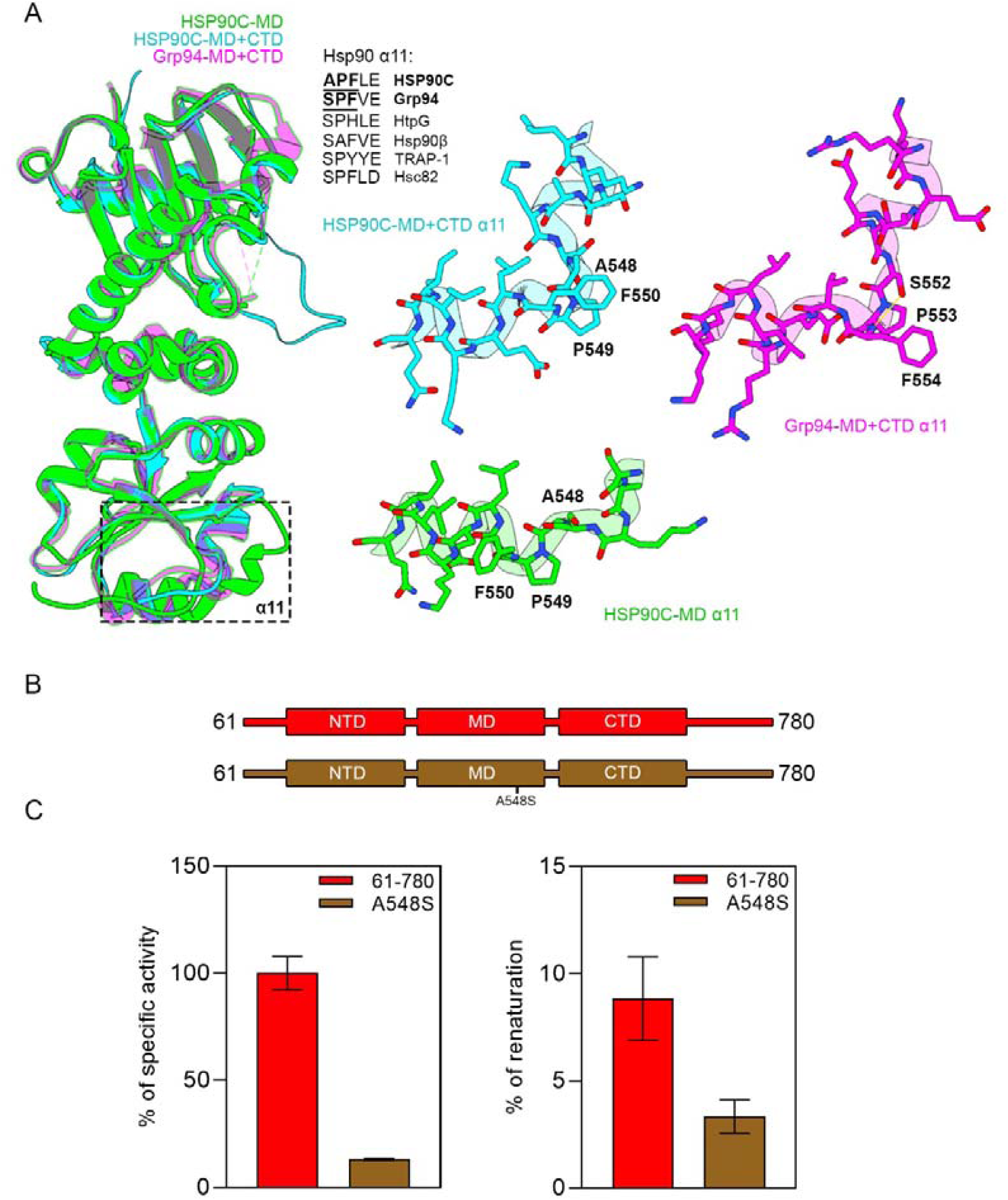
The α11 helical switch and its importance for HSP90C function. A: Superimposition of HSP90C middle domain (green) (PDB: 9SX3), HSP90C MD-CTD dimer (cyan) (PDB: 9SWT), and Grp94 MD-CTD (purple) (PDB: 2O1T). The α11 adopts different conformations in both HSP90C structures, due to a canonical serine among HSP90 proteins replaced by an alanine. B: Sequences of mature HSP90C (red) and punctual mutant A548S (brown). C: Impact of the A548S mutation on HSP90C activities. Both ATPase and refolding activities are decreased for the mutant.

To assess the importance of these 2 conformations, we cloned and produced a punctual mutant of the A548 of HSP90C (A548S) (Fig. 5B). By reverting the A548 into a serine, we expect the hydrogen bond to occur, thereby stabilizing the curved form of the α11 helix of HSP90C instead of the straight conformation. We then performed ATPase and refolding activity assays on this mutant to evaluate the impact of the mutation (Fig. 5C). Both of the activities were dramatically lower for the mutant in comparison to the wild-type, which highlights the importance of the A548 residue. Thermal shift assays indicated that the stability of HSP90C was not impacted by the mutation (Fig. S3). These last experiments show that the A548 residue is mandatory for the correct function of HSP90C.

Lastly, we determined kinetic parameters of both mature HSP90C (61-780) and A548S punctual mutant (Table 1, Fig. S4). This analysis revealed that the A548 mutation had no significant impact on ATP affinity, but decreased its hydrolysis rate (k_cat_) by 2-fold. Moreover, the best fitted-kinetic model was cooperative for 61-780, whereas the A548S mutant follows a Michaelis-Menten-type kinetic model.

**Table 1:** Kinetics parameters of mature WT HSP90C (61-780) and the A548S punctual mutant.

| HSP90C | Best model | $K_m$ (mM) | $K_{half}$ (mM) | $k_{cat}$ (min<sup>-1</sup>) | n | R<sup>2</sup> |
| --- | --- | --- | --- | --- | --- | --- |
| 61-780 | Hill (cooperative) | - | 1.61 ± 0.18 | 6.75 ± 0.01 | 2.02 ± 0.40 | 0.989 |
| A548S | Michaelis-Menten | 1.46 ± 0.19 | - | 3.26 ± 0.00 | 1 | 0.994 |

Taken with the crystallographic data, we propose that HSP90C undergoes a conformational switch through its ATPase cycle that involves a curved to straight movement of the α11 helix.

### Features of HSP90C among green plants

Lastly, we wanted to evaluate the significance of three of the mechanisms that we unveiled (the redox regulation in the N-terminal cap, the CTD extension and the conformational switch) among plants. Indeed, these three mechanisms are characterized by specific HSP90C residues: C61 for the N-terminal cap regulation, A548 for the helical switch, and R740-W741 for the CTD extension. For this purpose, we aligned *A. thaliana*’s HSP90C sequence against 14 representative species from green plants (*Viridiplantae*) (Fig. 6A). Subsequently, every best match from each species was selected, and used for multiple sequence alignment. The results revealed that without surprise, the three main domains of HSP90C are largely conserved among green plants, as well as for the loop between the middle domain and the C-terminal domain (Fig. S5). On the other hand, the most variable parts are the transit peptide, the N-terminal cap, the loop between the N-terminal domain and the middle domain, and the C-terminal tail.

**Figure 6:**
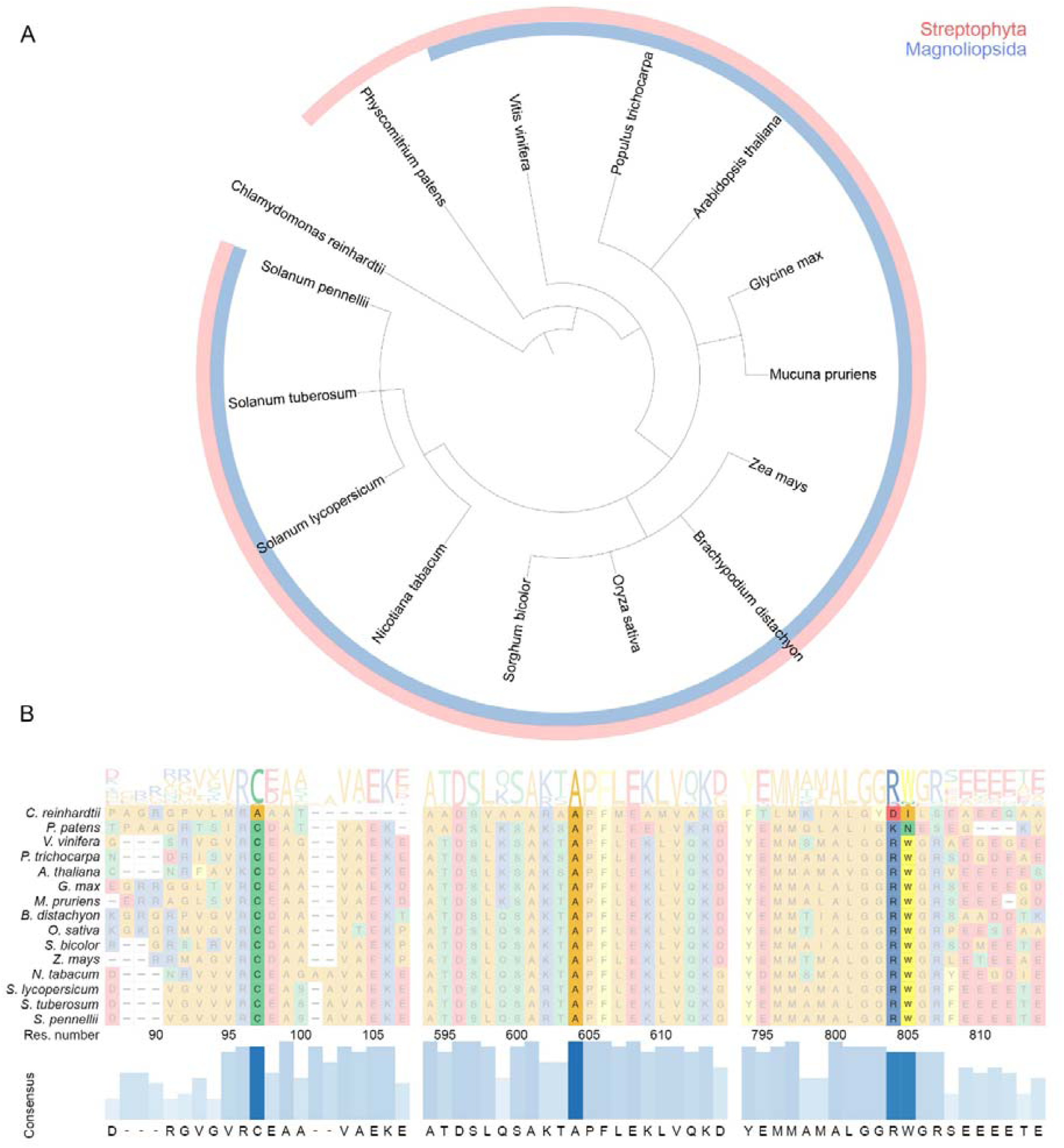
Conservation of HSP90C signature sequences among green plants. A: The phylogenetic tree features 14 species from green plants (*Viridiplantae*), including *C. reinhardtii* (algae), represented as two groups: Streptophyta and Magnoliopsida. B: Sequence conservation of C61 (1), A548 (2) and R740-W741 (3) from *A. thaliana’s* HSP90C in all represented species. Logo sequences are shown on top, as well as with the consensus sequence on the bottom.

Then, we specifically looked at the three HSP90C’s sequence signatures that we described in our work: C61, A548 and R740-W741 (Fig. 6B). First, the C61 residue is greatly conserved among green plants, except for *Chlamydomonas reinhardtii* (green algae). In this species, an alanine substitutes both this position and the next one. Furthermore, in all other species of green plants selected here (which are denominated as Streptophyta), the cysteine is followed by an acidic residue (D or E). Second, the A548 residue is perfectly conserved among all green plants represented here, including *Chlamydomonas reinhardtii*. Lastly, we evaluated the conservation of the two amino-acids constituting the extended dimeric interface in *A. thaliana*’s HSP90C (R740 and W741). The alignment revealed that only two groups of green plants do not contain this motif (*Chlamydomonas reinhardtii* and *Physcomitrium patens*). The subgroup of green plants that share this motif are classified as Magnoliopsida.

This analysis suggests that all the sequence signatures of *A. thaliana*’s HSP90C that we described here are shared by the Magnoliopsida class among green plants.

## Discussion

Throughout this study, we aimed to provide a molecular view of HSP90C, which has been previously described as crucial for plant integrity. The ATPase activity assays that we conducted in a first instance raised more questions, as HSP90C appeared to cycle very fast in comparison to the other HSP90 family members. Here we pinpoint atypical mechanisms occurring in HSP90C that explain its high activity and so its important role in the chloroplast.

First, we investigated the N-terminal cap of HSP90C. Our experiments showed that the C61 is involved in an intermonomeric disulfide bond, which stabilizes the dimer. We propose that this bond is a C61-C61 bond that connects the HSP90C monomers. Indeed, we demonstrated that C165 is not responsible for this bond, and C61 is the only remaining partner in range of such an interaction according to the AlphaFold3 prediction. The ATPase activity tests that we conducted revealed a lower activity when this bond is reduced. In parallel, the refolding activity assays indicated that HSP90C can refold client proteins *in vitro* more efficiently when the bond is reduced. One possible explanation of both these observations is that the C61 disulfide bond is limiting the opening of the dimer by bridging the 2 NTD of HSP90C monomers. Consequently, HSP90C can cycle faster in the presence of this bond due to a faster transition between open and closed states. However, this limited opening of the dimer may hinder some surfaces interacting with clients, thereby decreasing the foldase activity of HSP90C. Nevertheless, we here used a human refolding system (hHsp70 and DNAJB1 proteins) and a model client protein (the luciferase) to study the foldase activity of HSP90C *in vitro*. Therefore, it is possible that HSP90C interacts *in vivo* with client proteins from the chloroplast through different surfaces. Considering this, the C61 disulfide bond may regulate the HSP90C foldase activity differently *in vivo* than in our *in vitro* experiments. Besides, what is particularly relevant is the redox regulation of this bond by H_2_O_2_. Our experiments showed that in the presence of a reducing agent, the ATPase activity of HSP90C is lower, while its refolding activity increases. Upon H_2_O_2_ addition that follows the reduction, HSP90C is able to recover both its ATPase activity and foldase properties observed after its reduction. Considering that H_2_O_2_ is one of the products of photosynthesis that accumulates onto the chloroplast ^21–23^, this could be a regulation mechanism occurring in plants, thereby allowing HSP90C to have a specific behavior when there are various levels of ROS in the chloroplast.

Second, we studied the C-terminal tail of HSP90C. A recently published study showed that the end of the C-terminal tail (750-780) is required for efficient ATPase activity and client binding (Mu et al., 2024). Here we showed that a RWGRVE sequence located at the very start of the tail (739-745) is mandatory for the dimerization of HSP90C, which explains the loss of both ATPase and refolding activities that we observed. Further investigation with crystallography experiments revealed that R740 and D686 form a salt bridge, while W741 reinforces the dimeric hydrophobic interface, which both stabilizes the dimer. Usually, the HSP90 proteins dimerize through their CTD helical bundle. Furthermore, some HSP90 homologs like HtpG or TRAP-1 do not have any C-terminal tail after their CTD, and are still able to dimerize properly. Here, we demonstrated that the beginning of the C-tail of HSP90C plays an essential role in the dimerization process, by providing an extension of the helical bundle of the CTD through its R740-W741 motif. However, the need of such an interaction to maintain the dimeric state raises questions, to which we found hypothetical answers by further investigating the structures.

By superimposing the available structures of MD and CTD dimers of HSP90 homologs with our structure of the 333-745 HSP90C dimer, we observed both a wide opening of the dimer and a shift of secondary structures occurring in the CTD. As the NTD is not present in any of these structures, the MD position appears to be either tighter or wider positioned in the case of the full closed or opened state of Grp94, Hsc82, and Hsp90α. Thus, it is believed that the MD of HSP90C significantly moves upon opening and closing of the dimer. However, the CTD is usually not greatly affected by this conformational transition, as it is the anchor point of the HSP90 dimerization. The HSP90C dimer’s center helices (650-670) constitute the main lacking dimer interface among the HSP90 members that we analyzed. In a previously described structure of Grp94, a mimetic client peptide has been found located above these helices ^18^. In HSP90C, a new empty cavity is accessible due to the space between the center helices. Therefore, it may have important implications in client binding, which may possibly dock until the top of the CTD helix bundle. Besides, the shift of CTD’s secondary structures may also explain the importance of the RWGRVE sequence at the start of the C-terminal tail. Because of a wider opening, HSP90C may need additional interactions to keep its dimeric form upon the ATPase cycle, which involves the intermonomeric R740-D686 salt bridge and the extended hydrophobic interface involving W741.

Lately, we solved the structure of HSP90C’s middle domain (333-598), and superimposed it with our structure of the MD-CTD dimer (333-745). This revealed that the α11 helix of HSP90C can adopt two conformations: a “curved” and a “straight” one. While the “curved” conformation has been observed in all the published structures of HSP90 homologs, the “straight” conformation is unique and characteristic from HSP90C. This straight conformation is most likely to be due to the absence of a hydrogen bond from the sidechain of the serine that usually stabilizes the two parts of the α11 helix. As the replacing alanine is unable to make this hydrogen bond, the α11 helix is more flexible in the case of HSP90C, which may allow a switch between these two conformations. We demonstrated that by substituting A548 with a serine, both ATPase and refolding activities of HSP90C were greatly reduced. Therefore, the α11 helix is likely to be a key region undergoing conformational changes upon the ATPase cycle of HSP90C. This switch may also have important implications on the HSP90C’s dimer global shape, which could explain the high distance observed between the center helices described above. Consequently, HSP90C may need additional interactions to maintain its dimeric conformation, as for instance the CTD extension motif (R740-W741).

Lastly, we performed a multiple sequence alignment using HSP90C sequences from 14 representative species of green plants. The alignment showed that C61, A548 and the R740-W741 CTD motif that we identified for *A. thaliana* are perfectly conserved through the representatives of the Magnoliopsida class (13 species here). This suggests that the mechanisms related to these residues that we described here for *A. thaliana* are likely to occur in these species, which emphasizes the relevance of these findings. Besides, a larger class of green plants represented here (named Streptophyta) share the cysteine located in the N-terminal cap of HSP90C. This class includes all Magnoliopsida as well as *Physcomitrium patens*. Thus, the cysteine from the N-terminal cap may also regulate HSP90C function in these species. Regarding the evolution, this mechanism may had appeared before the CTD extension, as it is shared by less-evolved plants. Finally, the alanine responsible for the conformational switch that we identified in *A. thaliana* (A548) is conserved in all green plants represented here, including *Chlamydomonas reinhardtii*. Therefore, this signature could be the first one that appeared in HSP90C through evolution, suggesting that all HSP90C represented here may share the same atypical ATPase cycle. Besides, this analysis enriches our hypothesis of the relation between the conformational switch and the need of the CTD extension. Indeed, through evolution, the alanine signature appeared prior to the arginine-tryptophane motif. Thus, this last event may had occurred to compensate the dimer instability induced by the alanine mutation, thereby increasing both the stability and the yield of HSP90C.

## Conclusion

HSP90C has been shown to be crucial for plant growth and thylakoid formation. It is associated with proteins from both the chloroplast and the thylakoid membranes. Moreover, HSP90C participates in the assembly of PSII, as it potentiates the maturation of PsbO1, one of the PSII subunits. Using various biophysical approaches, we provide a molecular view of HSP90C. We identified an intermonomeric disulfide bond occurring in the N-terminal cap of HSP90C that involves C61. This bond can regulate both ATPase and refolding activities of HSP90C, and is sensitive to H_2_O_2_. On the other hand, we showed that the start of the C-terminal tail of HSP90C is required for its correct dimerization. More specifically, a 6 amino-acid CTD extension (RWGRVE) is connecting the two HSP90C monomers via a salt-bridge (R740-D686) and a reinforced dimeric interface (W741). Our crystal structures of HSP90C molecular constructions revealed a wide opening of the dimer, as well as a large shift of secondary structures in the CTD compared to its canonical position in HSP90 dimers from other species. Furthermore, we pinpointed an alternative conformation of the α11 helix, which is due to the presence of a conserved alanine residue. The mutation of this alanine greatly decreased both ATPase and foldase activities of HSP90C, which shows the importance of this particular amino-acid. We propose that the α11 helix of HSP90C undergoes a conformational change that is required for its activity, which possibly explains the repositioning of the CTD in the dimer and the extended dimeric interface. All these features are likely to be shared by most of representative of green plants, which includes *C. reinhardtii* regarding the helical switch. These findings provide explanations of the high ATPase activity of HSP90C among the HSP90 family, and may help for the understanding of client proteins folding in the chloroplast.

## Methods

### Cloning of HSP90C constructions

For the truncated forms, the HSP90C gene was amplified by PCR from *A. Thaliana* cDNA using the following primers: 5’-ATTAGCCATATGTGTGACGCCGCCGTGG-3’ for position 61, 5’-ATTAGCCATATGGTGGCGGAGAAAGAGACC-3’ for the position 65, 5’-CTTAGCCATATGGGGTCAGGTGAGAAGTTTG-3’ for the position 74, 5’-CTTAGCCATATGAAACCGCTATGGATGCGCAAT-3’ for the position 333 (forward), 5’-TAGTGCTCGAGTCAATCTTCATCTCCGAGTTCCAAA-3’ for the position 598, 5’-TAGTGCTCGAGTCATCCTCCAACCGCCATTG-3’ for the position 739, 5’-TAGTGCTCGAGTCATTCAACTCTGCCCCATCTTC-3’ for the position 745 and 5’-TAGTGCTCGAGTCAATCTTGCCAAGGATCACT-3’ for the position 780 (reverse). Each PCR product was inserted into a pET28a plasmid that contains a 6xHis peptide and a TEV cleavage site upstream the MCS using NdeI/XhoI restriction enzymes. *E.Coli* bacteria from NEB were transformed with the recombinant plasmid and selected on LB-kanamycin plates. For the punctual mutants (A548S and C165S), a directed mutagenesis kit was used to generate the mutated plasmids from the recombinant plasmid of the 61-780 construct. The primers for the C165S mutant were the following ones: 5’-CCTTGGAACTATTGCTCAAAGTG-3’ (forward) and 5’-GAATCAATAAGTTCTTCCTTTGTCATTC-3’ (reverse). For the A548S mutant, we used the following primers: 5’-TGCCAAGTCTAGCCCTTTCTTG-3’ (forward) and 5’-CTTTTAAGACTATCAGTTGC-3’ (reverse). Eventually, the plasmids were extracted and inserted into Rosetta cells for protein expression.

### Protein expression and purification

All the constructions were expressed into Rosetta cells from NEB in a 2YT medium at 37°C 180 rpm until DO at 600 nm reached 0.8. IPTG was then added to the medium at a concentration of 0.5 mM, and temperature set to 18°C for overnight growing. Pellets were collected after 4400g centrifugation at 4°C during 35 min, and freezed at 4°C for one day. After thawing, the pellets were resuspended in lysis buffer (25 mM Tris.HCl pH 8.0; 500 mM NaCl; 20 mM imidazole pH 8.0; supplemented with SIGMAFAST™ antiprotease (Sigma) and DNAse), and solubilized in cold room (4°C) for 30 minutes under agitation. Cells were disrupted by French press, with 2 runs at a pressure of 16,000 PSI (1.1 kbar) and then centrifugate at 15,000 g for 20 minutes. After filtering through 0.22 µm sized-pore filters, supernatants were injected into a NiNTA affinity column on Äkta GO system (Cytiva), and then equilibrated with non-supplemented lysis buffer. Elution was performed with 8 CV linear gradient from 20 M to 500 mM of imidazole. Fractions corresponding to the major eluted peak were pooled and diluted at 50 mM NaCl, in order to reduce salt concentration. Then, the sample was injected into a Q Sepharose Fast Flow ion-exchange column (Cytiva), equilibrated with washing buffer (25 mM Tris.HCl pH 8.0; 50 mM NaCl). After elution with a gradient from 50 mM to 500 mM of NaCl on 25 CV, the main peak was collected, and concentrated using a Amicon Ultra 10K (Merck Millipore). Lastly, the sample was filtered again with 0.22 µm sized-pore filters and injected into a Superdex 200 size-exclusion column (Cytiva), equilibrated with gel filtration buffer (25 mM Tris.HCl pH 8.0; 150 mM NaCl; 0.5 mM EDTA). Eventually, the fractions corresponding to the main peak were pooled, concentrated as before, and froze in liquid nitrogen before storing at -80°C.

### ATPase activity assays

The steady-state ATPase activity of Heat Shock Proteins was assessed using a coupled enzymatic assay (PK/LDH) that tracks NADH oxidation, as previously described ^24^, with a modified reaction buffer (50 mM HEPES pH 7.5; 100 mM KCl; 5 mM ATP; 5 mM MgCl₂). Reactions were carried out in 96-well plates and recorded on a CLARIOstar reader (BMG Labtech). HSP90 proteins were used at a concentration of 0.5 μM. To account for background ATPase activity, measurements for HSP90 were corrected by subtracting the signal obtained after the addition of 50 μM radicicol.

### Refolding activity assays

The ability of HSP90C to refold clients was assessed with a human *in vitro* chaperoning system, constituted by hHsp70, DNAJB1 and denaturated QuantiLum Recombinant Luciferase (Promega) as described before ^25^. Luciferase was diluted to 55 μM in denaturation buffer containing 6 M guanidium/HCl and 1 mM DTT during 30 min at room temperature. Denatured luciferase was diluted 125 times (0.4 μM) in renaturation buffer (20 mM Tris-HCl pH 7.5, 50 mM KCl, 5 mM MgCl_2_, 1 mM DTT) supplemented with 10 µM of hHsp70, 2 µM of DNAJB1 and 2 µM of Hsp90. Refolded luciferase activity was assessed by mixing 5 μL aliquots of samples with 120 μL of luciferase buffer (20 mM Tris-HCl, pH 7.5; 200 μM luciferin; 0.5 mM ATP; 10 mM MgCl₂). Chemiluminescence was recorded using a CLARIOstar plate reader (BMG Labtech). The percentage of refolded luciferase was determined relative to the activity of native luciferase.

### SDS-PAGE

The SDS-PAGE gel was poured with a 12.5% stacking solution and a 5% resolving solution. Samples were mixed with laemmli buffer. In reducing conditions, the denaturation mix contained 143 mM of 2-mercaptoethanol. The mixtures were heat at 95°C for 5 minutes, and loaded onto the gel. Migration was performed at 140 V for 70 min. The gel was stained with a Coomassie Blue solution for 60 min, and destained for 60 minutes before being transferred in milliQ water.

### SEC-MALS experiments

The SEC-MALS setup used compris a Shimadzu HPLC-UV system coupled with an Optilab T-rEX refractometer and a miniDawn TREOS MALS detector (Wyatt Technology). A sample volume of 20 µl at a concentration of 2.5 mg/ml was injected onto a Superose 6 Increase 10/300 GL column (Cytiva), which was equilibrated at a flow rate of 0.4 ml/min using the following buffer: 50 mM Tris (pH 7.5), 150 mM NaCl and 0.5 mM EDTA. Prior to the experiments, a run with a monodisperse BSA sample was performed under the same chromatographic conditions to calibrate the SEC-MALS (normalisation of the light scattering detectors, inter-detector delay correction, and band broadening correction). The concentration detector is the refractometer, with a value of 0.1850 ml/g used for the analyte refractive index increment. The weight-averaged molar mass of the analyte was determined using Astra 5.3.4 software (Wyatt Technology).

### Crystallogenesis

For the HSP90C 333-745 construction, first crystals were obtained using the sitting-drop method in plates with the JCSG Plus screen solution from Hampton Research screening. From the well G2, the crystallization mix was optimized to a final solution containing 20 mM MgCl_2_, 0.1 M HEPES pH 7.5, 28% polyacrylic acid, with the addition of 8.5% of ethylene glycol. The drop was made with 50 nL of a 30.2 mg/mL protein solution and 50 nL of the well solution. After 3 weeks, the crystals reached their final shape, and were collected using a CryoLoop from Hampton Research. No cryo protectant was added.

For the HSP90C 333-598 construction, initial crystals were obtained using the JCSG Plus screen from Hampton Research at the D7 position (0.2 M LiSO_4_, 0.1 M Tris pH 8.5, 40% PEG 400) with the sitting-drop method. The drop was made with 50 nL of a 32.2 mg/mL protein solution and 50 nL of the well solution. After 2 weeks, crystals were collected, with no addition of cryo protectant.

### Crystallography data collection and processing

The crystallography data was collected at Synchrotron SOLEIL in Saclay, France, using PROXIMA-1 (HSP90C 333-745) and PROXIMA-2 (HSP90C 333-598) beam lines. Images were processed using XDS ^26^, and the resulting HKL file was scaled using autoPROC ^27^. The output was used as an input for molecular replacement in the PHENIX suit ^28–31^. For the HSP90C dimer, the data were significantly anisotropic and were ellipsoidally truncated and scaled with STARANISO ^32^. The statistics for the data collection are shown in Table S1.

### Model building and refinement

For molecular replacement, we used Phaser-MR ^33^ from the PHENIX suit ^28–31^. Individual domains of Grp94 MD+CTD dimer from *C. lupus familiaris* ^34^ (PDB: 2O1T) and HtpG middle domain from *E. coli* ^35^ (PDB: 2GQ0) were used as initial models. For refinement, we used phenix.refine to perform a first iteration with rigid body fit enabled. From here, successive rounds of manual check on Coot ^36,37^ and refinement in PHENIX were conducted, with non-crystallographic symmetries and grouped-ADP enabled (all other options set to default). All the refinement statistics are shown in Table S1.

### Phylogenetic tree

The phylogenetic tree was built using the Taxonomy Browser tool from NCBI ^38^. Identically as a previous study, the following 14 species were selected: *Chlamydomonas reinhardtii*, *Physcomitrium patens*, *Vitis vinifera*, *Populus trichocarpa*, *Arabidopsis thaliana*, *Glycine max*, *Mucuna pruriens*, *Brachypodium distachyon*, *Oryza sativa*, *Sorghum bicolor*, *Zea mays*, *Nicotiana tabacum*, *Solanum lycopersicum*, *Solanum tuberosum*, and *Solanum pennellii*. The tree was visualized using interactive Tree Of Life (iTOL) ^39^, with default parameters and branch lengths ignored.

### Multiple sequence alignments

All multiple sequence alignments were outputted by Clustal Omega from EMBL-EBI ^40^, with default parameters. For alignments that were used in the Figure 6, the output was visualized with ggmsa package ^41^ using R.

## Acknowledgments

We acknowledge the Synchrotron SOLEIL (Gif-sur-Yvette, France) for provision of synchrotron radiation facilities and thank the beamline staff of PROXIMA-1 and PROXIMA-2A for their assistance during data collection. This project received financial support from the ANR LABEX DYNAMO (ANR-11-LABX 0011).

---

**Figure S1:**
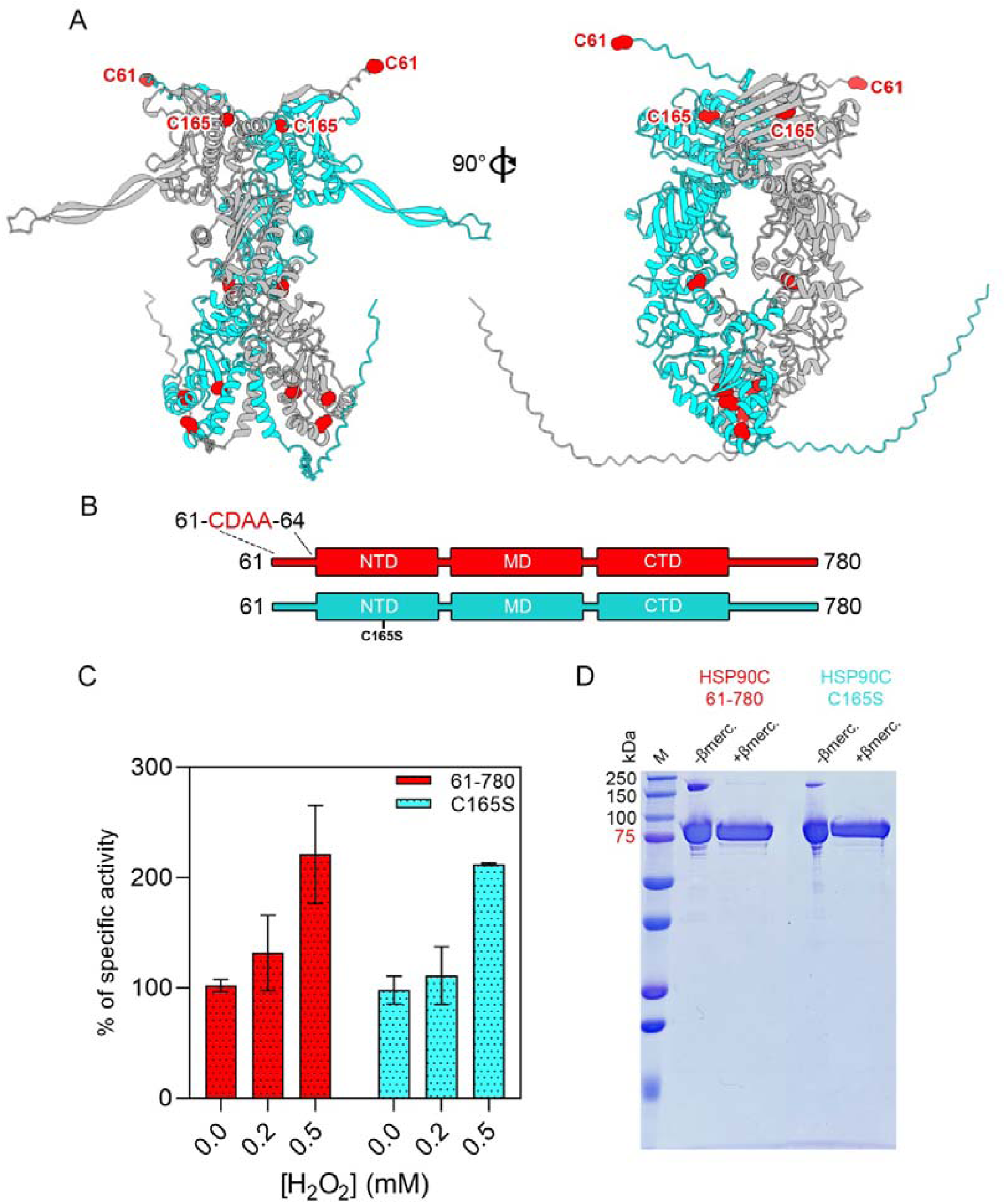
Analysis of the intermonomeric disulfide bridge of the N-terminal cap. A: AlphaFold3 dimer prediction, with all HSP90C’s cysteines shown as red spheres. B: Sequences of mature HSP90C 61-780 (red) and C165S punctual mutant (cyan). C: ATPase activity recovering in the presence of H_2_O_2_. D: SDS-PAGE of HSP90C 61-780 and C165S, showing a dimeric band in non-reducing conditions only.

**Figure S2:**
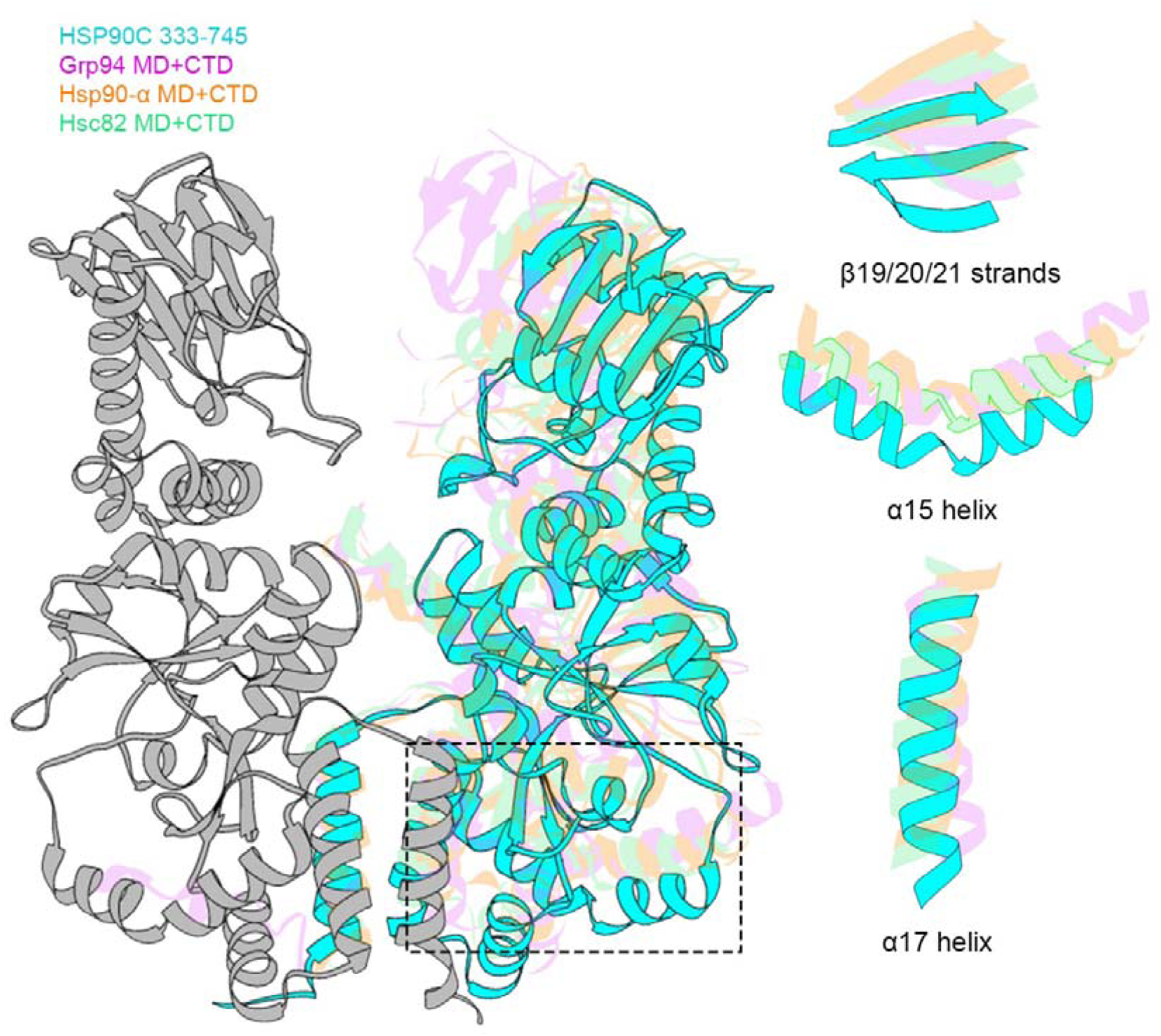
Superimposition of HSP90 MD-CTD dimers from different species. Dimers includes only the middle domains and C-terminal domains of HSP90C from *A. Thaliana* (PDB: 9SWT) (cyan), Grp94 from *C. lupus familiaris* (PDB: 2O1T) (purple), Hsc82 from *S. cerevisiae* (PDB: 2CGE) (light green), and Hsp90α from *H. sapiens* (PDB: 7RY1) (orange). All HSP90 dimers were aligned on the CTD of one monomer (only the HSP90C monomer is shown here in grey), to appreciate the relative opening of each dimer. The shifted CTD secondary structures are zoomed on the right panel.

**Figure S3:**
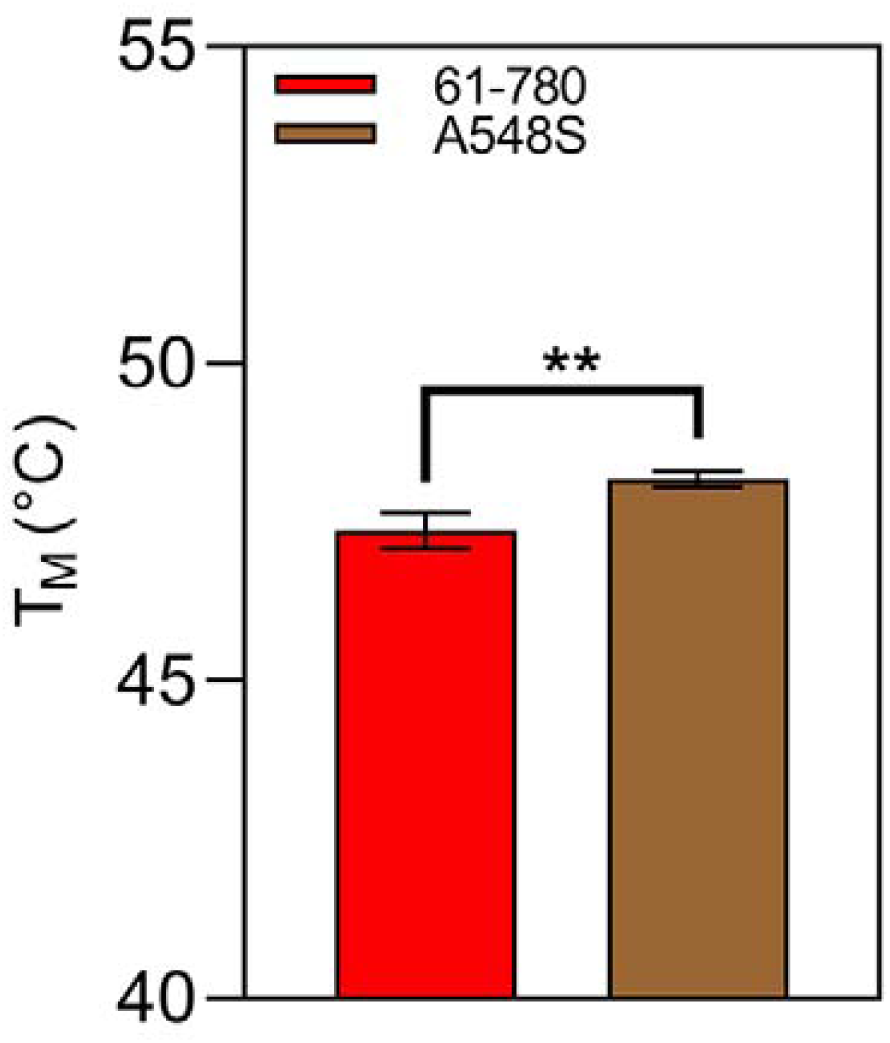
Thermal Shift Assay (TSA) of mature HSP90C (61-780) and the A548S punctual mutant. Both constructions have similar T_M_, which indicates that the mutation did not affect dramatically the stability of the mutant.

**Figure S4:**
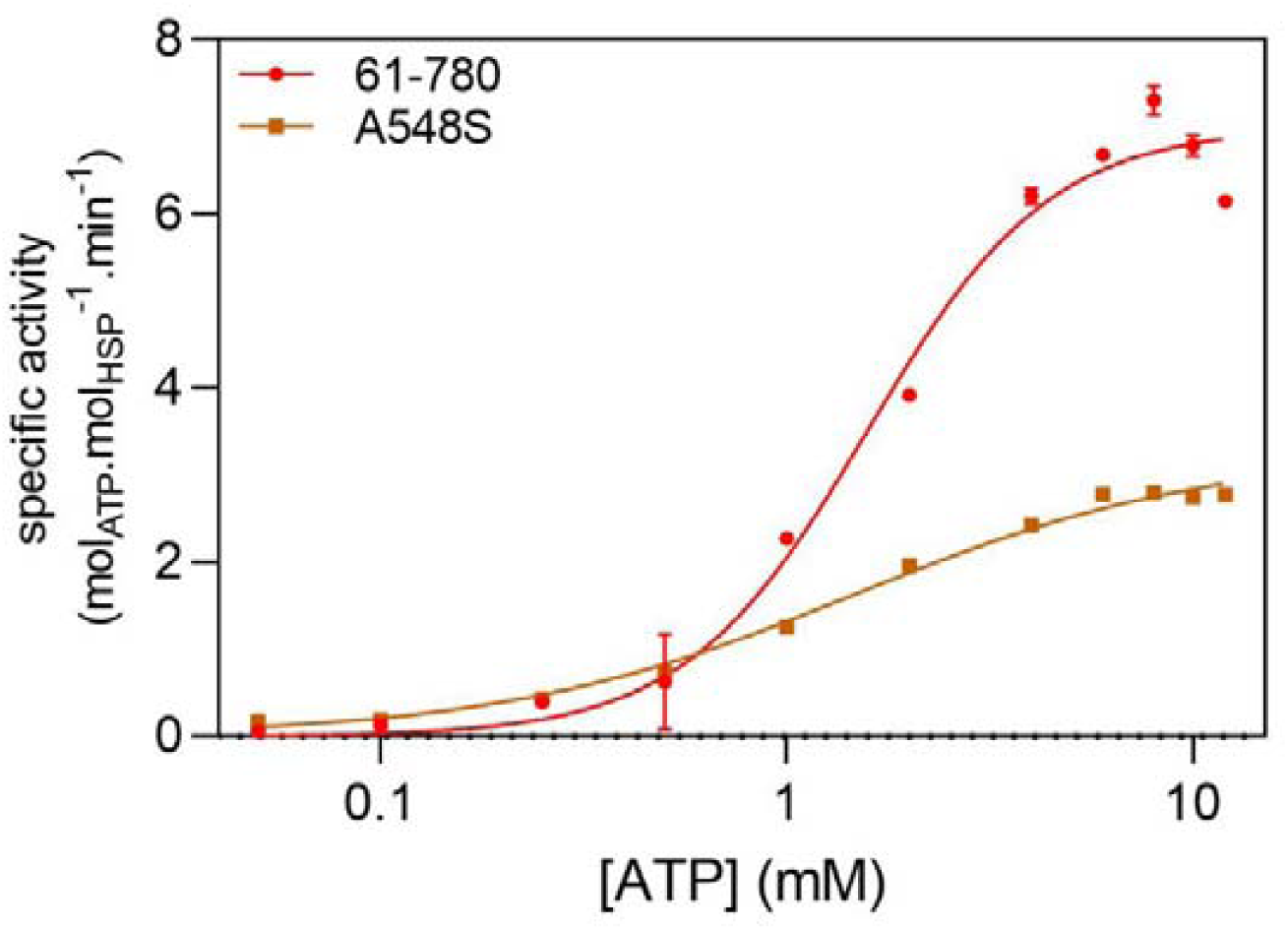
Curve-fitting of specific activity measurements of mature HSP90C (61-780) and the A548S punctual mutant. HSP90C 61-780 (red) follows a cooperative model (Hill), while A548S (brown) fits a Michaelis-Menten model. Parameters shown in Table 1 were calculated from this fit.

**Figure S5:**
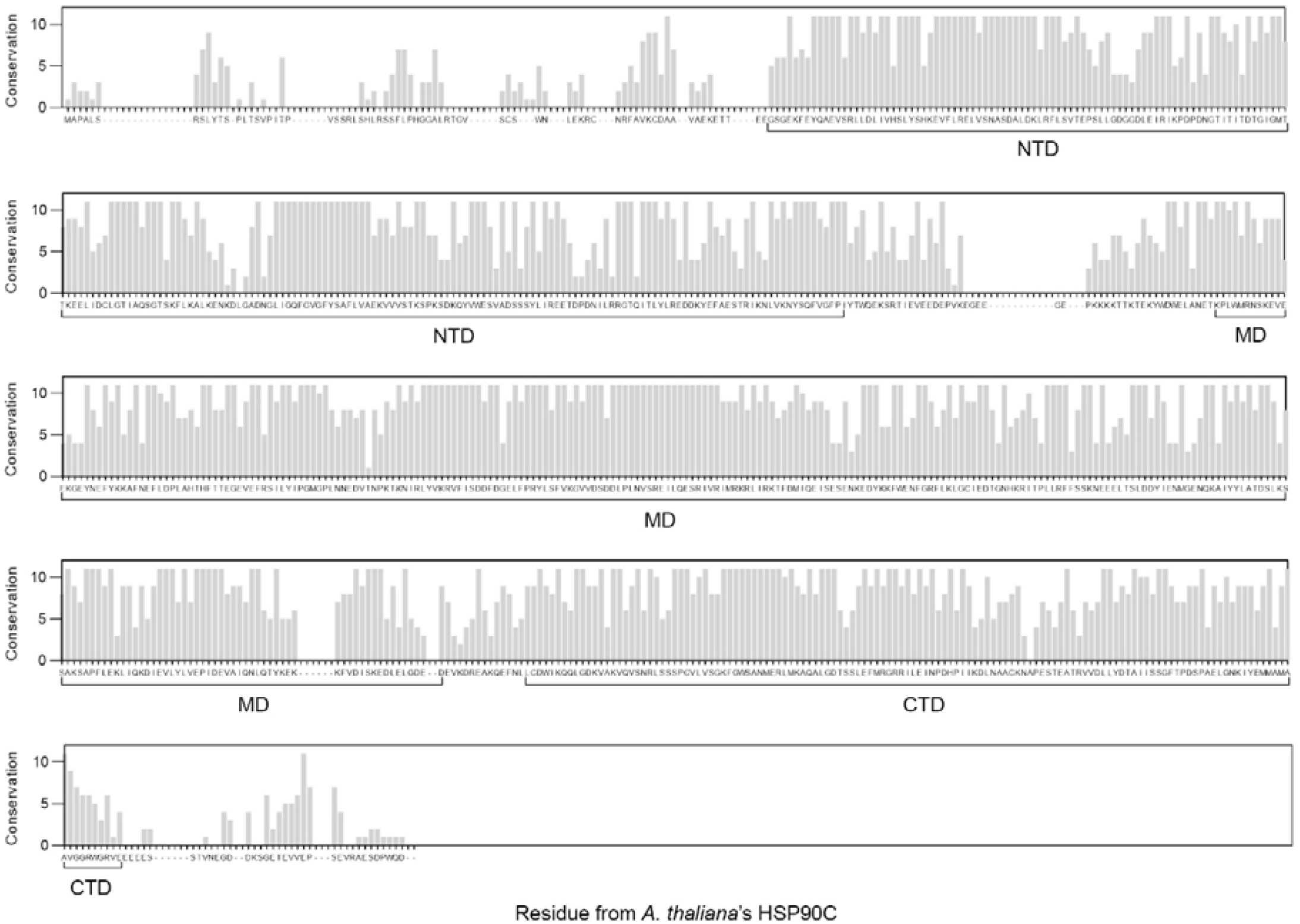
Conservation of the whole *A. thaliana*’s HSP90C sequence among representative of green plants. The following 14 species were selected: *Chlamydomonas reinhardtii*, *Physcomitrium patens*, *Vitis vinifera*, *Populus trichocarpa*, *Arabidopsis thaliana*, *Glycine max*, *Mucuna pruriens*, *Brachypodium distachyon*, *Oryza sativa*, *Sorghum bicolor*, *Zea mays*, *Nicotiana tabacum*, *Solanum lycopersicum*, *Solanum tuberosum*, and *Solanum pennellii*.

**Table S1:** Crystallographic data and refinement statistics.

|  | Middle domain of HSP90C | Middle and C-terminal domains of HSP90C (dimer) |
| --- | --- | --- |
| PDB entry | 9SX3 | 9SWT |
| **Data collection** |  |  |
| Wavelength | 0.9801 | 0.9786 |
| Resolution range | 70.72 - 2.92 (2.97 - 2.92) | 53.06 - 3.147 (3.6 - 3.15) |
| Space group | C 1 2 1 | C 2 2 21 |
| Unit cell | 132.01 84.03 60.37 90 95.88 90 | 82.28 170.62 135.53 90 90 90 |
| Total reflections | 79246 (4134) | 235265 (7692) |
| Unique reflections | 14455 (727) | 9723 (486) |
| Multiplicity | 5.5 (5.7) | 24.2 (15.8) |
| Completeness (%) | 99.9 (100) | - |
| Completeness (spherical) (%) | - | 57.4 (9.4) |
| Completeness (ellipsoidal) (%) | - | 91.1 (67.0) |
| Mean I/sigma(I) | 13.2 | 14.2 |
| Wilson B-factor | 97.28 | 131.72 |
| R-merge | 0.086 | 0.127 |
| R-meas | 0.095 | 0.130 |
| R-pim | 0.040 | 0.026 |
| CC1/2 | 0.998 (0.658) | 1 (0.552) |
| **Refinement** |  |  |
| Resolution range in refinement | 47.09 - 3.01 (3.24 - 3.01) | 53.06 - 3.15 (3.60 - 3.15) |
| Reflections used in refinement | 12936 (2430) | 9712 (568) |
| Reflections used for R-free | 647 (122) | 478 (30) |
| R-work | 0.2029 (0.3747) | 0.2174 (0.3582) |
| R-free | 0.2585 (0.4569) | 0.2767 (0.4012) |
| Number of non-hydrogen atoms | 4308 | 6674 |
| Macromolecules | 4291 | 6666 |
| Ligands | 15 | 6 |
| Solvent | 2 | 2 |
| Protein residues | 515 | 822 |
| RMS (bonds) | 0.010 | 0.003 |
| RMS (angles) | 1.19 | 0.70 |
| Ramachandran favored (%) | 92.34 | 92.67 |
| Ramachandran allowed (%) | 5.70 | 6.72 |
| Ramachandran outliers (%) | 1.96 | 0.61 |
| Rotamer outliers (%) | 0.00 | 0.00 |
| Clashscore | 11.31 | 8.03 |
| Average B-factor | 105.60 | 141.81 |
| Macromolecules | 105.27 | 141.81 |
| Ligands | 200.20 | 143.28 |
| Solvent | 105.45 | 126.07 |
| MolProbity score | 2.04 | 1.90 |

## References

1. Dobrogojski, J., Adamiec, M. & Luciński, R. The chloroplast genome: a review. Acta Physiol. Plant 42, 98 (2020).

2. Jin, Z., Wan, L., Zhang, Y., Li, X., Cao, Y., Liu, H., Fan, S., Cao, D., Wang, Z., Li, X., Pan, J., Dong, M.-Q., Wu, J. & Yan, Z. Structure of a TOC-TIC supercomplex spanning two chloroplast envelope membranes. Cell 185, 4788–4800.e13 (2022).

3. Xu, X., Ouyang, M., Lu, D., Zheng, C. & Zhang, L. Protein Sorting within Chloroplasts. Trends Cell Biol. 31, 9–16 (2021).

4. Schmitz, G., Schmidt, M. & Feierabend, J. Characterization of a plastid-specific HSP90 homologue: identification of a cDNA sequence, phylogenetic descendence and analysis of its mRNA and protein expression. Plant Mol. Biol. 30, 479–492 (1996).

5. Cao, D., Lin, Y. & Cheng, C. L. Genetic interactions between the chlorate-resistant mutant cr 8 8 and the photomorphogenic mutants cop1 and hy5. Plant Cell 12, 199–210 (2000).

6. Oh, S. E., Yeung, C., Babaei-Rad, R. & Zhao, R. Cosuppression of the chloroplast localized molecular chaperone HSP90.5 impairs plant development and chloroplast biogenesis in Arabidopsis. BMC Res. Notes 7, 643 (2014).

7. Inoue, H., Li, M. & Schnell, D. J. An essential role for chloroplast heat shock protein 90 (Hsp90C) in protein import into chloroplasts. Proc. Natl. Acad. Sci. U. S. A. 110, 3173–3178 (2013).

8. Willmund, F. & Schroda, M. HEAT SHOCK PROTEIN 90C is a bona fide Hsp90 that interacts with plastidic HSP70B in Chlamydomonas reinhardtii. Plant Physiol. 138, 2310–2322 (2005).

9. Jiang, T., Oh, E. S., Bonea, D. & Zhao, R. HSP90C interacts with PsbO1 and facilitates its thylakoid distribution from chloroplast stroma in Arabidopsis. PLoS One 12, e0190168 (2017).

10. Jiang, T., Mu, B. & Zhao, R. Plastid chaperone HSP90C guides precursor proteins to the SEC translocase for thylakoid transport. J. Exp. Bot. 71, 7073–7087 (2020).

11. Heide, H., Nordhues, A., Drepper, F., Nick, S., Schulz-Raffelt, M., Haehnel, W. & Schroda, M. Application of quantitative immunoprecipitation combined with knockdown and cross-linking to Chlamydomonas reveals the presence of vesicle-inducing protein in plastids 1 in a common complex with chloroplast HSP90C. Proteomics 9, 3079–3089 (2009).

12. Mu, B., Nair, A. M. & Zhao, R. Plastid HSP90C C-terminal extension region plays a regulatory role in chaperone activity and client binding. Plant J. 119, 2288–2302 (2024).

13. Hoter, A., El-Sabban, M. E. & Naim, H. Y. The HSP90 Family: Structure, Regulation, Function, and Implications in Health and Disease. Int. J. Mol. Sci. 19, (2018).

14. Panaretou, B., Prodromou, C., Roe, S. M., O’Brien, R., Ladbury, J. E., Piper, P. W. & Pearl, L. H. ATP binding and hydrolysis are essential to the function of the Hsp90 molecular chaperone in vivo. EMBO J. 17, 4829–4836 (1998).

15. Meyer, P., Prodromou, C., Hu, B., Vaughan, C., Roe, S. M., Panaretou, B., Piper, P. W. & Pearl, L. H. Structural and functional analysis of the middle segment of hsp90: implications for ATP hydrolysis and client protein and cochaperone interactions. Mol. Cell 11, 647–658 (2003).

16. Huai, Q., Wang, H., Liu, Y., Kim, H.-Y., Toft, D. & Ke, H. Structures of the N-terminal and middle domains of E. coli Hsp90 and conformation changes upon ADP binding. Structure 13, 579–590 (2005).

17. Pearl, L. H. & Prodromou, C. Structure and mechanism of the Hsp90 molecular chaperone machinery. Annu. Rev. Biochem. 75, 271–294 (2006).

18. Huck, J. D., Que, N. L., Hong, F., Li, Z. & Gewirth, D. T. Structural and Functional Analysis of GRP94 in the Closed State Reveals an Essential Role for the Pre-N Domain and a Potential Client-Binding Site. Cell Rep. 20, 2800–2809 (2017).

19. Lavery, L. A., Partridge, J. R., Ramelot, T. A., Elnatan, D., Kennedy, M. A. & Agard, D. A. Structural asymmetry in the closed state of mitochondrial Hsp90 (TRAP1) supports a two-step ATP hydrolysis mechanism. Mol. Cell 53, 330–343 (2014).

20. Marzec, M., Eletto, D. & Argon, Y. GRP94: An HSP90-like protein specialized for protein folding and quality control in the endoplasmic reticulum. Biochim. Biophys. Acta 1823, 774–787 (2012).

21. Slesak, I., Libik, M., Karpinska, B., Karpinski, S. & Miszalski, Z. The role of hydrogen peroxide in regulation of plant metabolism and cellular signalling in response to environmental stresses. Acta Biochim. Pol. 54, 39–50 (2007).

22. Fahnenstich, H., Scarpeci, T. E., Valle, E. M., Flügge, U.-I. & Maurino, V. G. Generation of hydrogen peroxide in chloroplasts of Arabidopsis overexpressing glycolate oxidase as an inducible system to study oxidative stress. Plant Physiol. 148, 719–729 (2008).

23. Breeze, E. & Mullineaux, P. M. The Passage of H2O2 from Chloroplasts to Their Associated Nucleus during Retrograde Signalling: Reflections on the Role of the Nuclear Envelope. Plants 11, (2022).

24. Henri, J., Chagot, M.-E., Bourguet, M., Abel, Y., Terral, G., Maurizy, C., Aigueperse, C., Georgescauld, F., Vandermoere, F., Saint-Fort, R., Behm-Ansmant, I., Charpentier, B., Pradet-Balade, B., Verheggen, C., Bertrand, E., Meyer, P., Cianférani, S., Manival, X. & Quinternet, M. Deep structural analysis of RPAP3 and PIH1D1, two components of the HSP90 co-chaperone R2TP complex. Structure 26, 1196–1209.e8 (2018).

25. Ramos, R., Karaiskou, A., Botuha, C., Amhaz, S., Trichet, M., Dingli, F., Forté, J., Lam, F., Canette, A., Chaumeton, C., Salome, M., Chenuel, T., Bergonzi, C., Meyer, P., Bohic, S., Loew, D., Salmain, M. & Sobczak-Thépot, J. Identification of cellular protein targets of a half-sandwich iridium(III) complex reveals its dual mechanism of action via both electrophilic and oxidative stresses. J. Med. Chem. 67, 6189–6206 (2024).

26. Kabsch, W. XDS. Acta Crystallogr. D Biol. Crystallogr. 66, 125–132 (2010).

27. Vonrhein, C., Flensburg, C., Keller, P., Fogh, R., Sharff, A., Tickle, I. J. & Bricogne, G. Advanced exploitation of unmerged reflection data during processing and refinement with autoPROC and BUSTER. Acta Crystallogr. D Struct. Biol. 80, 148–158 (2024).

28. Afonine, P. V., Grosse-Kunstleve, R. W., Urzhumtsev, A. & Adams, P. D. Automatic multiple-zone rigid-body refinement with a large convergence radius. J. Appl. Crystallogr. 42, 607–615 (2009).

29. Afonine, P. V., Grosse-Kunstleve, R. W., Echols, N., Headd, J. J., Moriarty, N. W., Mustyakimov, M., Terwilliger, T. C., Urzhumtsev, A., Zwart, P. H. & Adams, P. D. Towards automated crystallographic structure refinement with phenix.refine. Acta Crystallogr. D Biol. Crystallogr. 68, 352–367 (2012).

30. Headd, J. J., Echols, N., Afonine, P. V., Grosse-Kunstleve, R. W., Chen, V. B., Moriarty, N. W., Richardson, D. C., Richardson, J. S. & Adams, P. D. Use of knowledge-based restraints in phenix.refine to improve macromolecular refinement at low resolution. Acta Crystallogr. D Biol. Crystallogr. 68, 381–390 (2012).

31. Afonine, P. V., Grosse-Kunstleve, R. W., Adams, P. D. & Urzhumtsev, A. Bulk-solvent and overall scaling revisited: faster calculations, improved results. Acta Crystallogr. D Biol. Crystallogr. 69, 625–634 (2013).

32. Tickle, I.J., Flensburg, C., Keller, P., Paciorek, W., Sharff, A., Vonrhein, C., Bricogne, G. STARANISO anisotropy & Bayesian estimation server. (2016). at <https://staraniso.globalphasing.org/cgi-bin/staraniso.cgi>

33. McCoy, A. J., Grosse-Kunstleve, R. W., Adams, P. D., Winn, M. D., Storoni, L. C. & Read, R. J. Phaser crystallographic software. J. Appl. Crystallogr. 40, 658–674 (2007).

34. Dollins, D. E., Warren, J. J., Immormino, R. M. & Gewirth, D. T. Structures of GRP94-nucleotide complexes reveal mechanistic differences between the hsp90 chaperones. Mol. Cell 28, 41–56 (2007).

35. Shiau, A. K., Harris, S. F., Southworth, D. R. & Agard, D. A. Structural Analysis of E. coli hsp90 reveals dramatic nucleotide-dependent conformational rearrangements. Cell 127, 329–340 (2006).

36. Emsley, P. & Cowtan, K. Coot: model-building tools for molecular graphics. Acta Crystallogr. D Biol. Crystallogr. 60, 2126–2132 (2004).

37. Emsley, P., Lohkamp, B., Scott, W. G. & Cowtan, K. Features and development of coot. Acta Crystallogr. D Biol. Crystallogr. 66, 486–501 (2010).

38. Schoch, C. L., Ciufo, S., Domrachev, M., Hotton, C. L., Kannan, S., Khovanskaya, R., Leipe, D., Mcveigh, R., O’Neill, K., Robbertse, B., Sharma, S., Soussov, V., Sullivan, J. P., Sun, L., Turner, S. & Karsch-Mizrachi, I. NCBI Taxonomy: a comprehensive update on curation, resources and tools. Database (Oxford) 2020, (2020).

39. Letunic, I. & Bork, P. Interactive Tree of Life (iTOL) v6: recent updates to the phylogenetic tree display and annotation tool. Nucleic Acids Res. 52, W78–W82 (2024).

40. Madeira, F., Madhusoodanan, N., Lee, J., Eusebi, A., Niewielska, A., Tivey, A. R. N., Lopez, R. & Butcher, S. The EMBL-EBI Job Dispatcher sequence analysis tools framework in 2024. Nucleic Acids Res. 52, W521–W525 (2024).

41. Zhou, L., Feng, T., Xu, S., Gao, F., Lam, T. T., Wang, Q., Wu, T., Huang, H., Zhan, L., Li, L., Guan, Y., Dai, Z. & Yu, G. Ggmsa: A visual exploration tool for multiple sequence alignment and associated data. Brief. Bioinform. 23, bbac222 (2022).

42. Robert, X. & Gouet, P. Deciphering key features in protein structures with the new ENDscript server. Nucleic Acids Res. 42, W320–4 (2014).

