## Supplementary figures and images for "Structural basis of HSP90C, a highly active chloroplastic HSP90 chaperone from *A. thaliana*"

### Figure S1

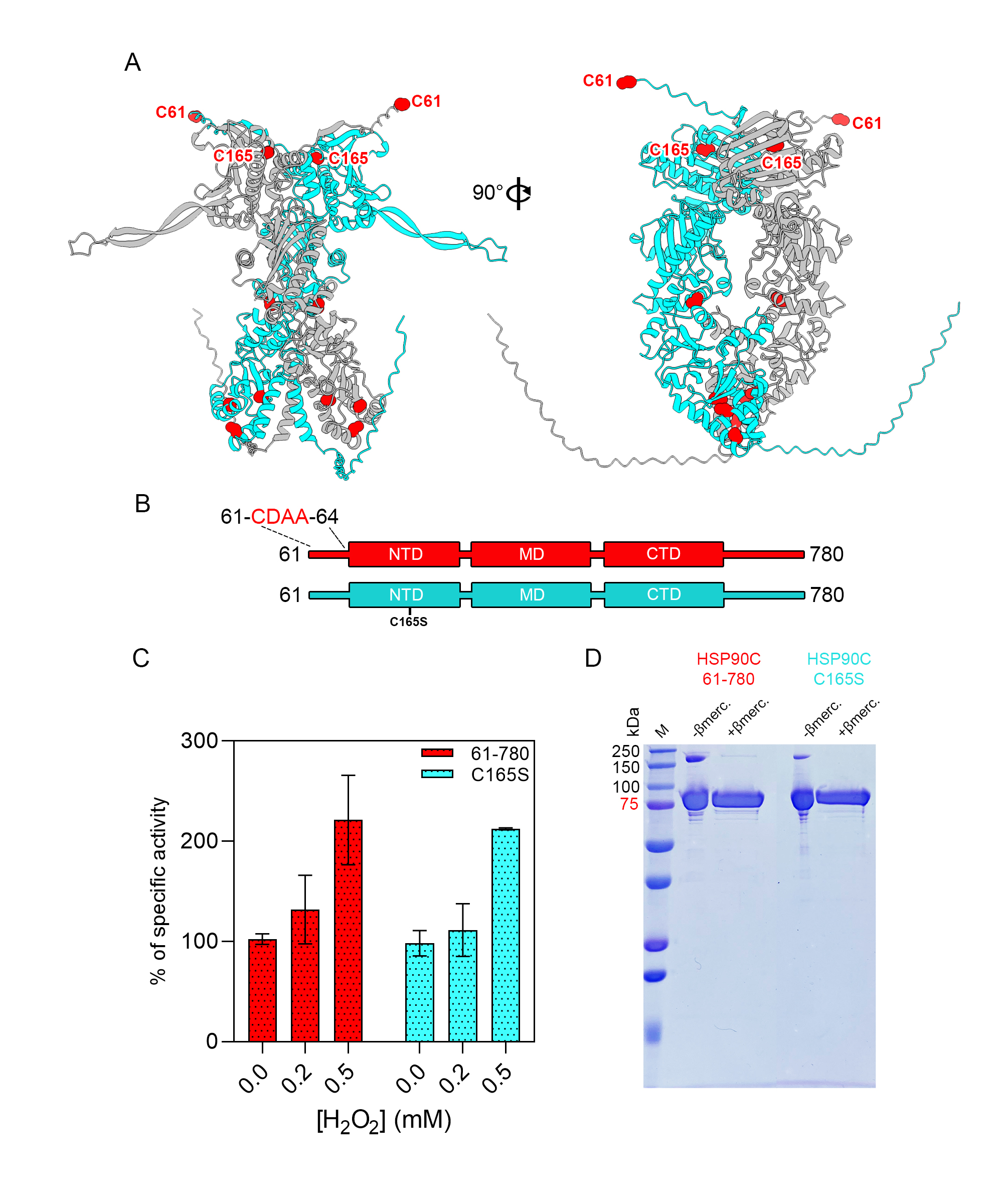

### Figure S2

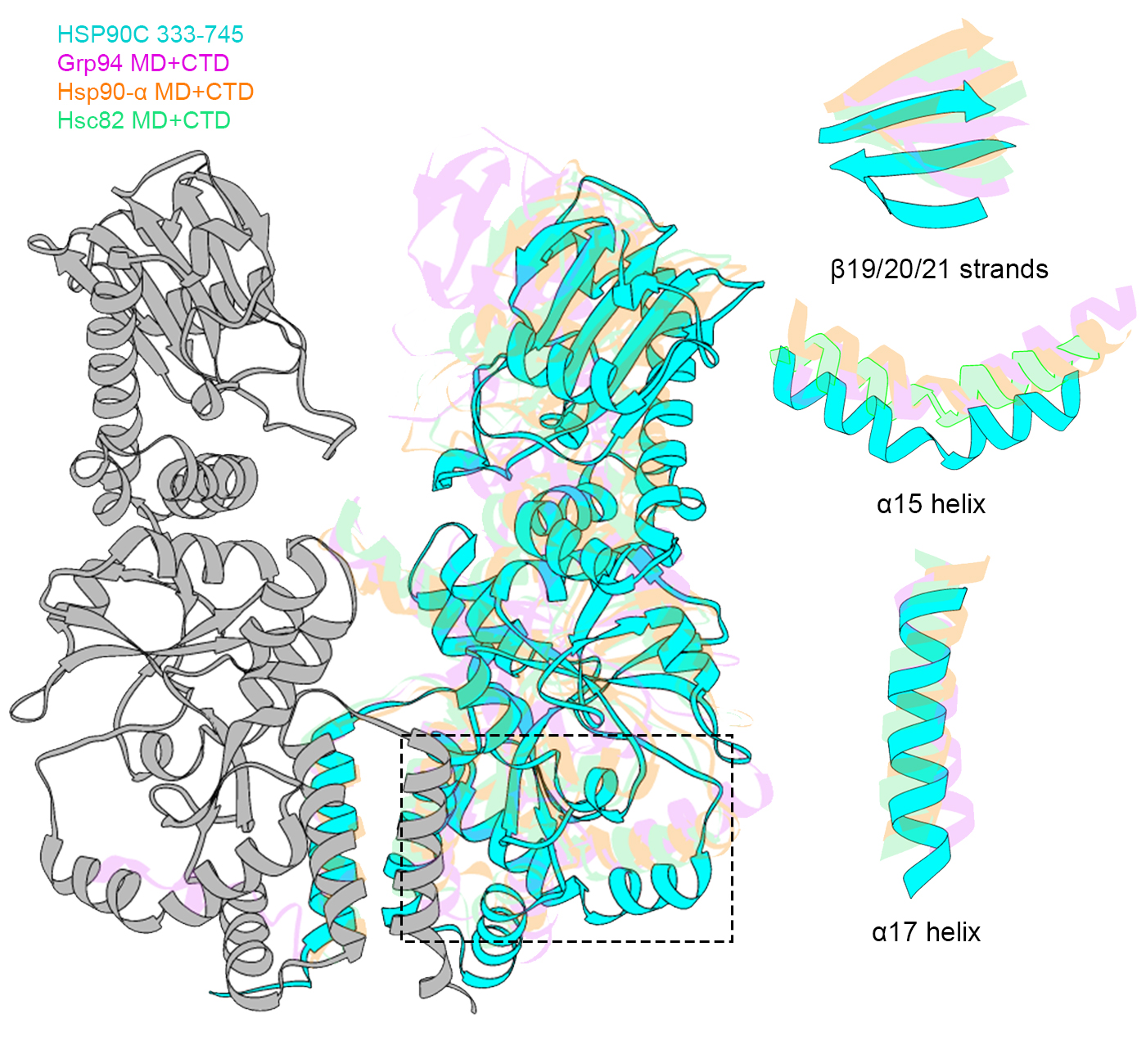

### Figure S3

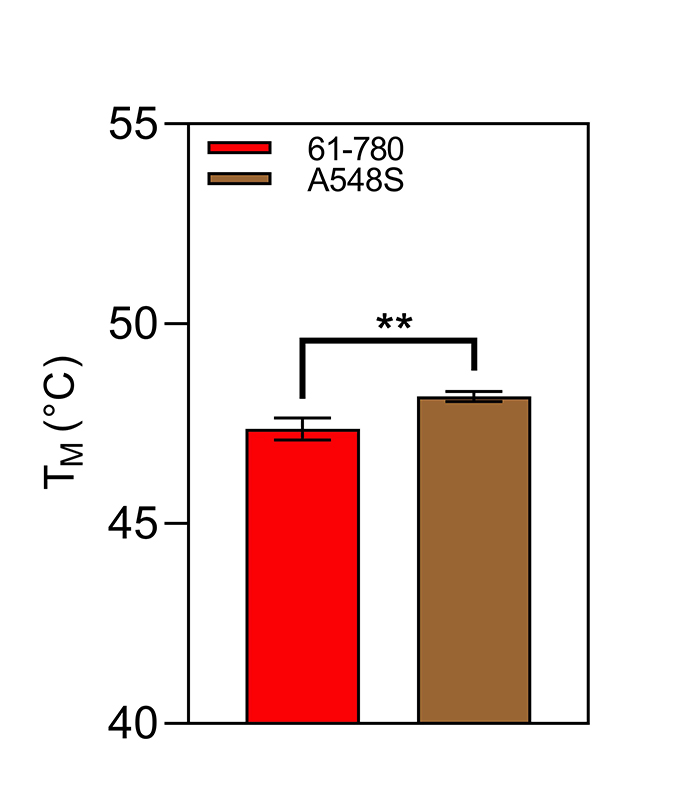

### Figure S4

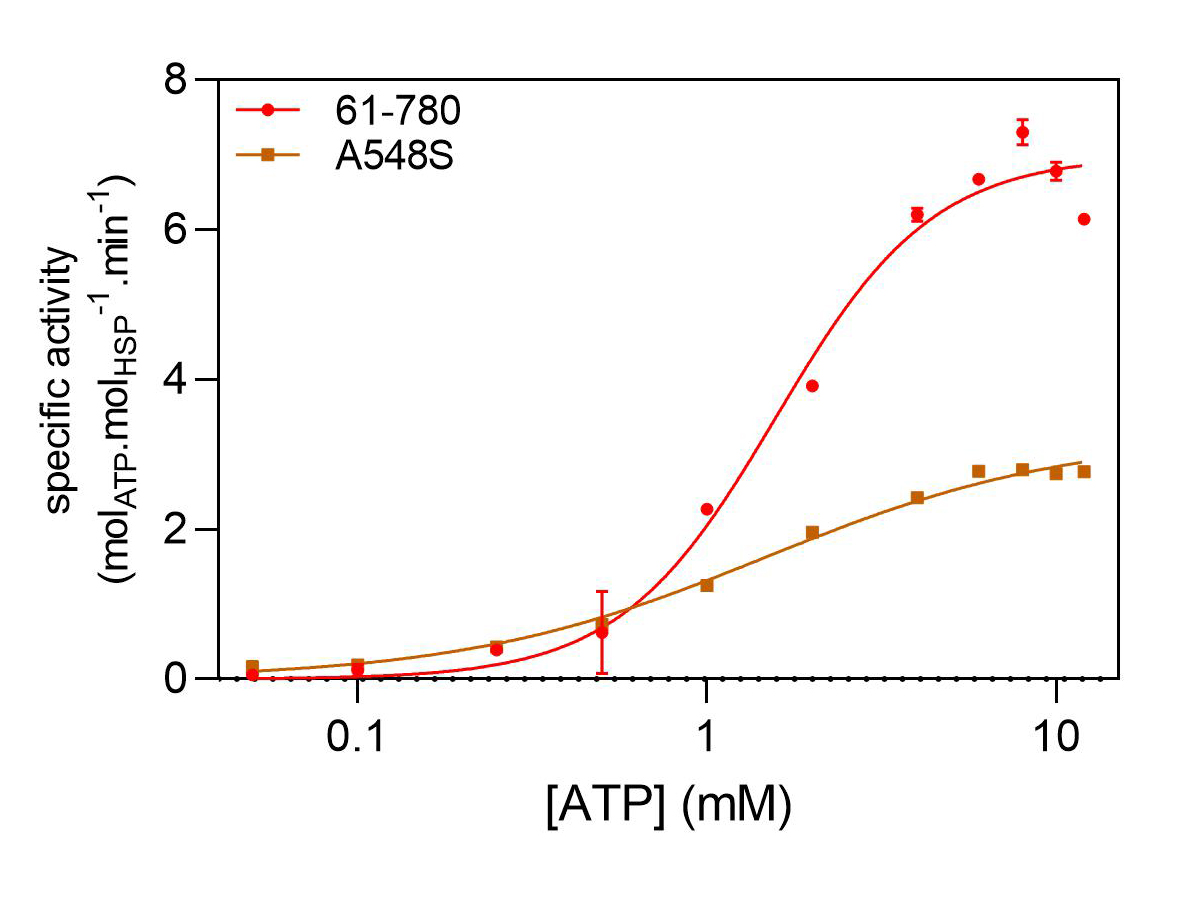

### Figure S5

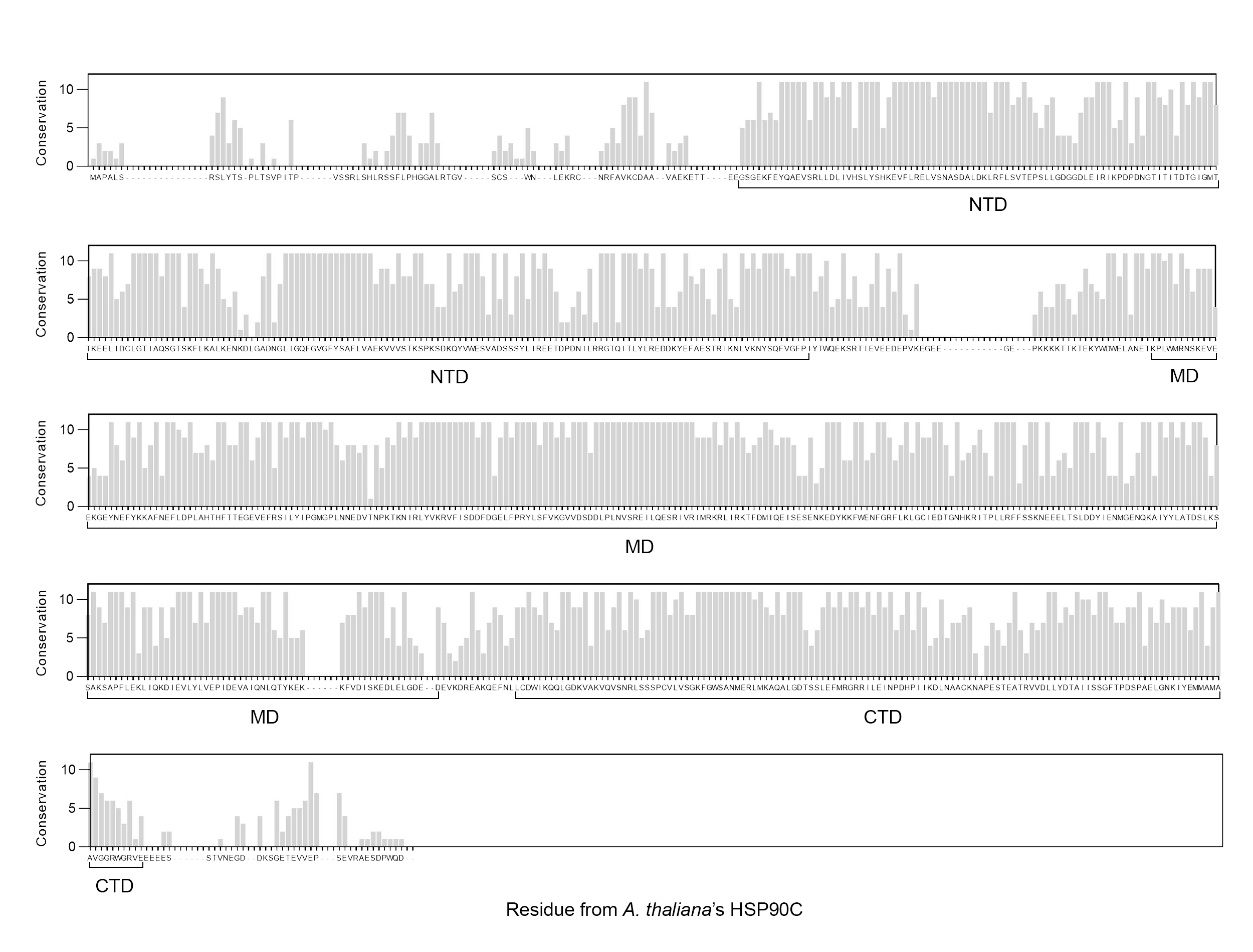
